# Multiomics profiling of zebrafish embryonic cell line PAC2 across growth phases to assess its relevance for toxicological studies

**DOI:** 10.1101/2025.07.26.666973

**Authors:** Mihai-Ovidiu Degeratu, Jessica Bertoli, Nikolai Huwa, David Lopez Rodriguez, Marion Revel, René Schönenberger, Colette vom Berg, Kristin Schirmer, Ksenia J. Groh

## Abstract

Permanent fish cell lines offer promising alternatives to traditional animal models for environmental risk assessment of chemicals. However, to facilitate their broader uptake into toxicity testing practice, a better understanding of functional capacities and expression of toxicologically relevant molecular targets is needed. Here, we present an extensive molecular profiling of the zebrafish embryonic cell line PAC2, combining global proteomics across cell population growth phases (over 7300 protein groups) with matched transcriptomics at exponential and stationary phase (over 14500 transcripts). Proteome coverage was sufficiently deep to reveal functional insights consistent with those derived from transcriptomics data, despite differences in the total number of measured genes. Major gene expression shifts detected upon transition from exponential to stationary cell population growth phase indicated reduction in DNA replication, translation, metabolism, and cell cycle regulation, along with increased stress responses, immune system responses, and extracellular matrix remodeling. Functional annotation revealed expression of core cellular processes along with a number of toxicologically relevant pathways, including xenobiotic metabolism, stress signaling, and nuclear receptors responsive to important chemical classes, such as steroids (e.g., estrogens, glucocorticoids) and chemicals known to disrupt lipid metabolism, e.g., through interaction with peroxisome proliferation activating receptors. These findings reinforce the potential of PAC2 cells to offer a versatile *in vitro* model for studying fish cell biology and omics-enhanced exploration of chemical toxicity mechanisms, aided by the well-developed molecular annotation in zebrafish. Moreover, the analysis approaches developed in this work offer a blueprint for molecular baseline characterization of other fish cell lines. This work thus strengthens the mechanistic foundation supporting the use of fish cell lines as alternative models in aquatic toxicity testing and risk assessment.

## Introduction

Chemical toxicity testing traditionally relies on *in vivo* tests using animal models. Fish are widely used as test animals for environmental risk assessment (1) due to their ecological relevance and sensitivity to environmental contaminants known to accumulate in freshwater systems (2). However, whole-animal fish tests are both resource-intensive and ethically contentious. As millions of fish are sacrificed annually to test chemicals and water samples (3, 4), these tests are increasingly scrutinized in light of the 3Rs (replacement, reduction and refinement) principles (5), similarly to other animal-based test methods.

In the last two decades, the toxicology community has witnessed an ongoing shift toward increased reliance on alternative (non-animal) test approaches (also known as new approach methodologies or NAMs) that aim to reduce or eliminate animal use (6). This new paradigm relies on the mechanistic understanding of toxicity, postulating that chemical-induced molecular and cellular changes precede higher-level effects and hence could be used to predict whole-organism toxicity outcomes (7, 8). Among ecotoxicological NAMs, permanent fish cell lines offer a variety of practical approaches to explore molecular and cellular toxicity mechanisms (3).

A wide variety of fish cell lines, derived from different tissues and species, are available (9, 10) and have been increasingly used for aquatic toxicity testing (11). Among these, cell lines derived from rainbow trout have been used most often (12–16), including as a model system for the ISO Standard (17) and the OECD Test Guideline 249 which use the gill-derived RTgill-W1 cell line for prediction of acute fish toxicity (18). The fish invitrome framework for animal-free toxicity assessment further builds on this OECD-adopted test by introducing a suite of additional modules, e.g., for bioaccumulation or chemical effects on growth (19). These modules can include additional cell lines derived from other organs or species, along with endpoint-specific test methods and computational models. In this context, cell lines from zebrafish (*Danio rerio*) could offer a unique advantage for molecular investigations thanks to extensive genomic and bioinformatic resources already available for this species (20–22).

One limitation currently hindering a broader applicability of the fish invitrome approach stems from insufficient knowledge about the molecular and functional repertoire of fish cell lines. Understanding toxicity-relevant properties and functional capacities of different fish cells lines, i.e., which genes and pathways are expressed and regulated under various conditions, could help to further improve their utility for toxicity testing. The growth of cell populations in *in vitro* culture undergoes two main phases: the exponential phase, characterized by active proliferation, and the stationary phase, marked by a plateau in cell proliferation, often coinciding with the formation of a confluent cell monolayer (23). Both phases have been purposefully used for toxicity assessment. Acute toxicity effects are typically assessed with cells in confluent monolayers, corresponding to stationary phase (24), while effects on cell population growth, predictive of chemical impacts on fish growth *in vivo*, are tested in exponentially proliferating cells (25). The transition from exponential to stationary phase is accompanied by massive gene expression shifts (26), which could also influence the expression of pathways relevant to stress responses and toxicity. Yet, such growth phase-specific gene expression changes have not yet been examined in detail in fish cell models. Among the various molecular profiling methods, transcriptomics has been used most widely due to the maturity of technology and deep coverage. However, mRNA levels alone often do not reliably reflect protein abundance, especially under dynamic conditions (27–29). Therefore, multiomic profiling, combining proteomics and transcriptomics, could enable a more comprehensive understanding of gene expression changes across growth phases (30–32).

In this study, we conducted an in-depth molecular characterization of the permanent zebrafish cell line PAC2 (33, 34), sampled in different phases of cell population growth and extensively profiled on the protein level with label-free mass spectrometry-based global proteomics. We further complemented these data by transcriptomics analysis with RNAseq, performed at two matching time points in order to gain a more comprehensive view of gene expression profiles. This integrated approach allowed us to (i) reveal mRNA and protein expression changes across different phases of PAC2 cell population growth; (ii) assess the degree of correlation between transcriptomic and proteomic profiles measured in matched samples and compare the insights delivered by the two techniques; (iii) explore the functional capacity of the PAC2 cells, particularly with regard to toxicologically relevant genes and pathways expressed by these cells. This information is expected to increase the value of the PAC2 cell line and facilitate its broader use in toxicological testing, as well as support its integration into the fish invitrome framework (19), especially for omics-enhanced assessments.

## Results

### Growth of PAC2 cells in culture and sampling for omics analyses

PAC2 cell population growth was characterized in 96-well plates over 3 weeks (Fig. 1 and Fig. S1). In the exponential growth phase, cell population doubling time (estimated between days 3-13) was ca. 3.3 days. Although the cell numbers measured by Hoechst nuclear staining appeared to increase continuously throughout the whole observation period, metabolic activity started plateauing around day 11 post-seeding (Fig. 1). Membrane integrity measurements also indicated that exponential growth could be slowing down around day 11-13 (Fig. S1). Moreover, metabolic activity-based curves resembled the shape of a curve constructed based on the direct cell counts data that we collected earlier in 12-well plates with otherwise similar culturing conditions (35). Therefore, also considering that the Hoechst staining of the nucleus does not distinguish between live and dead cells, we chose to rely on the metabolic activity measurements for selecting the time points to collect samples for omics analyses. Specifically, we sampled PAC2 cells on day 3, 8 and 14 post-seeding, assuming that these timepoints correspond to the early exponential, mid-exponential, and stationary phases of cell population growth (Fig. 1). These samples comprised proteomics dataset 1 (PD1). On days 8 and 14 of the same experiment, we also collected matched samples for transcriptomics analysis, from which we derived a transcriptomics dataset (TD). The inclusion of transcriptomics analysis in our study allowed increasing the number of profiled genes and enabled a direct comparison between molecular insights provided by proteomics and transcriptomics data.

**Figure 1.**
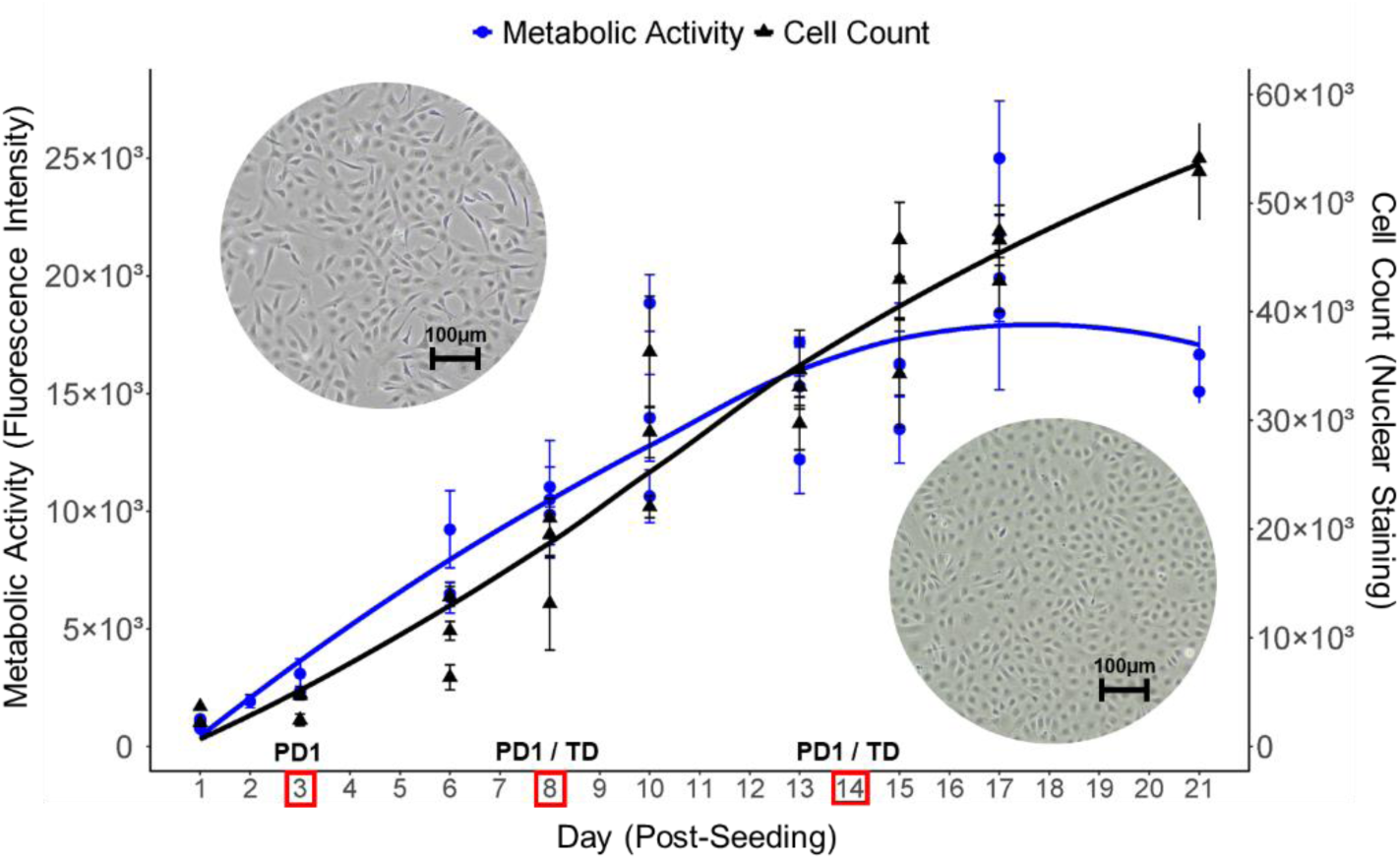
Growth curve of the PAC2 cell line based on nuclear staining counts and metabolic activity measurements. PAC2 cells were seeded in 96-well plates and cultured for three weeks. Cell culture medium was exchanged once on day 7 post-seeding. Metabolic activity based on alamarBlue fluorescence (arbitrary units, blue circles line, left y-axis) and cell counts based on Hoechst 33342 nuclear staining (black triangles line, right y-axis) were recorded in three biological replicates at multiple timepoints (days) throughout the culturing duration. Data are shown as mean ± SD of technical replicates (n=3-12) for each biological replicate. Trend lines were added using LOESS smoothing (geom_smooth, span = 1) to visualize overall changes over time. Red boxes and labels above them indicate the days on which the samples for proteomics dataset 1 (PD1) and transcriptomics dataset (TD) were collected. Representative phase contrast microscopy images included on the graph in the top left and bottom right corners show the appearance of PAC2 cells grown in T75 flasks at ca. 60% and 95% confluency, respectively, corresponding to exponential and stationary phases. Scale bar = 100µm.

In addition to PD1, we included in this study a second set of PAC2 samples, collected in the frame of a targeted phosphoproteomics study published previously (35), and used them to derive proteomics dataset 2 (PD2). These samples provided an even more resolved representation of different cell population growth phases: specifically, days 4, 7, 11, 18 and 28, assumed to correspond, respectively, to early, mid- and late exponential, and early and late stationary phases. The inclusion of the second proteomics dataset allowed us to enhance the depth and reliability of proteomics-derived insights into expression changes occurring across cell population growth phases. Further, while PD1 and PD2 samples differ in several parameters of experimental design (summarized in Table 1), they were analyzed by the same global proteomics data acquisition method in the frame of the current study. This additionally allowed us to address important technical questions related to the comparability of proteomics sample preparation procedures and robustness of mass spectrometry-based global proteomics analyses.

**Table 1.** Overview of experimental design parameters for omics datasets analyzed in this study.

| Experimental parameters | Transcriptomics dataset (TD) | Proteomics dataset 1 (PD1) | Proteomics dataset 2 (PD2) |
| --- | --- | --- | --- |
| PAC2 cell culture specifics | Plate format: 6-well plates<br>Media exchange: Day 7 post-seeding |  | Plate format: 12-well plates<br>Media exchange: Days 7, 14, 21 |
| Sampling points | Days 8, 14 | Days 3, 8, 14 | Days 4, 7, 11, 18, 28 |
| Sample preparation and storage | mRNA-enriched library, short storage at +4°C | Tryptic digest, non-enriched, desalted, short storage at +4°C | Tryptic digest, non-bound fraction from Fe-NTA phosphopeptide enrichment, longer-term storage at –70°C |
| Peptide loading in LC-MS/MS | Not relevant | 1 µg – Day 3<br>2 µg – Days 8, 14 | 4 µg – all timepoints |

### Proteomics profiling of PAC2 cells across cell population growth phases

Global proteomics analysis showed a high consistency of precursor, peptide, protein and protein group identification numbers across technical and biological replicates within both datasets, showing %CV values of 4.6%, 5.5%, 2.8% and 2.0%, respectively (see Supplementary File (SF) 1). Across both the PD1 (SF 2A) and PD2 (SF 3A) combined, up to 11796 proteins could be identified, corresponding to 7432 protein groups (protein groups comprise either uniquely identified proteins or sets of indistinguishable proteins inferred from the identified peptides). Among these, 112 represented contaminants and were therefore removed from downstream analysis. Over 95% (7165) of the identified 7320 *Danio rerio* protein groups were shared between the two datasets (Fig. 2, top Venn diagram). Similarly, high overlaps of the identified protein groups were observed between the different sampling days within each dataset (Fig. 2, bottom Venn diagrams). The abundance of measured proteins spanned over 6 orders of magnitude, with log10 median protein group quantity ranging from <1 to >7.

**Figure 2.**
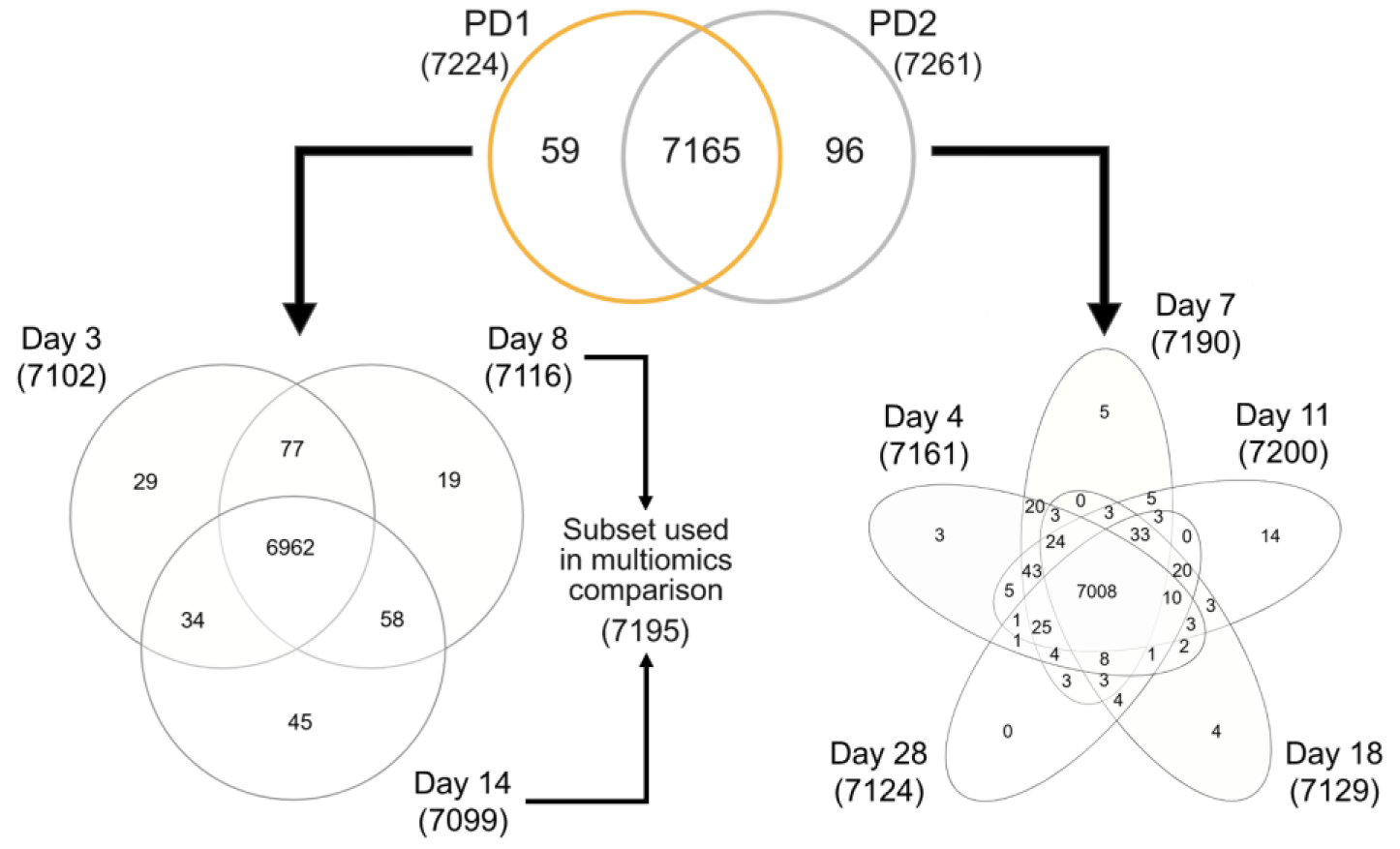
Numbers and overlap of protein groups identified within and between proteomics datasets 1 (PD1) and 2 (PD2). In total, 7320 protein groups were identified in the PAC2 cell line in the two datasets combined, the majority of which were shared between the two datasets (see top Venn diagram) and between the different sampling conditions (days) within the same dataset (see bottom Venn diagrams for PD1 (left) and PD2 (right)). 7195 PD1 protein groups detected in day 8 and day 14 samples were used for multiomics comparison with matched transcriptomics samples.

In the next step, identification data was prepared for quantitative analysis. For this, protein groups with <70% detection frequency were removed to ensure statistical power, and protein groups with detection frequency ≥70% were imputed if they did not appear in all replicates (see Fig. S2 for histograms of measured and imputed counts in each replicate). Following this procedure, the number of protein groups included in PD1 and PD2 was reduced to 6751 (SF 2B) and 6927 (SF 3B), respectively, and the combined list containing protein groups detected across PD1 and PD2 together was reduced to 6803 protein groups. Within these lists, less than 10% of protein groups were single hits (i.e., identified by only one peptide). For example, in the shared list of 6803 protein groups, 445 protein groups were single-peptide identifications. The reduced lists were used to perform quantitative analyses such as the principal component analysis (PCA) and differential protein expression analysis presented below.

PCA showed a clear separation between all conditions analyzed (Fig. 3A). Samples belonging to the two datasets (PD1 and PD2) were separated in principal component 1 (PC1), while PC2 primarily showed separation between growth phases (sampling days) within each dataset. A similar trend, i.e., a progression over time from early exponential to (late) stationary phase, was observed in both PD1 and PD2 (Fig. 3A). Specifically, days 3 and 8 from PD1 clustered closely together, similarly to days 4 and 7 from PD2. These samples, corresponding to exponential growth phase, were strongly separated from samples corresponding to the stationary growth phase, i.e., day 14 of PD1 and days 18 and 28 of PD2, the latter two closely clustered together too. Interestingly, day 11 of PD2 was located between the exponential- and stationary-phase related samples of this dataset, indicating that it could be reflective of a transition period between the two distinct growth phases.

**Figure 3.**
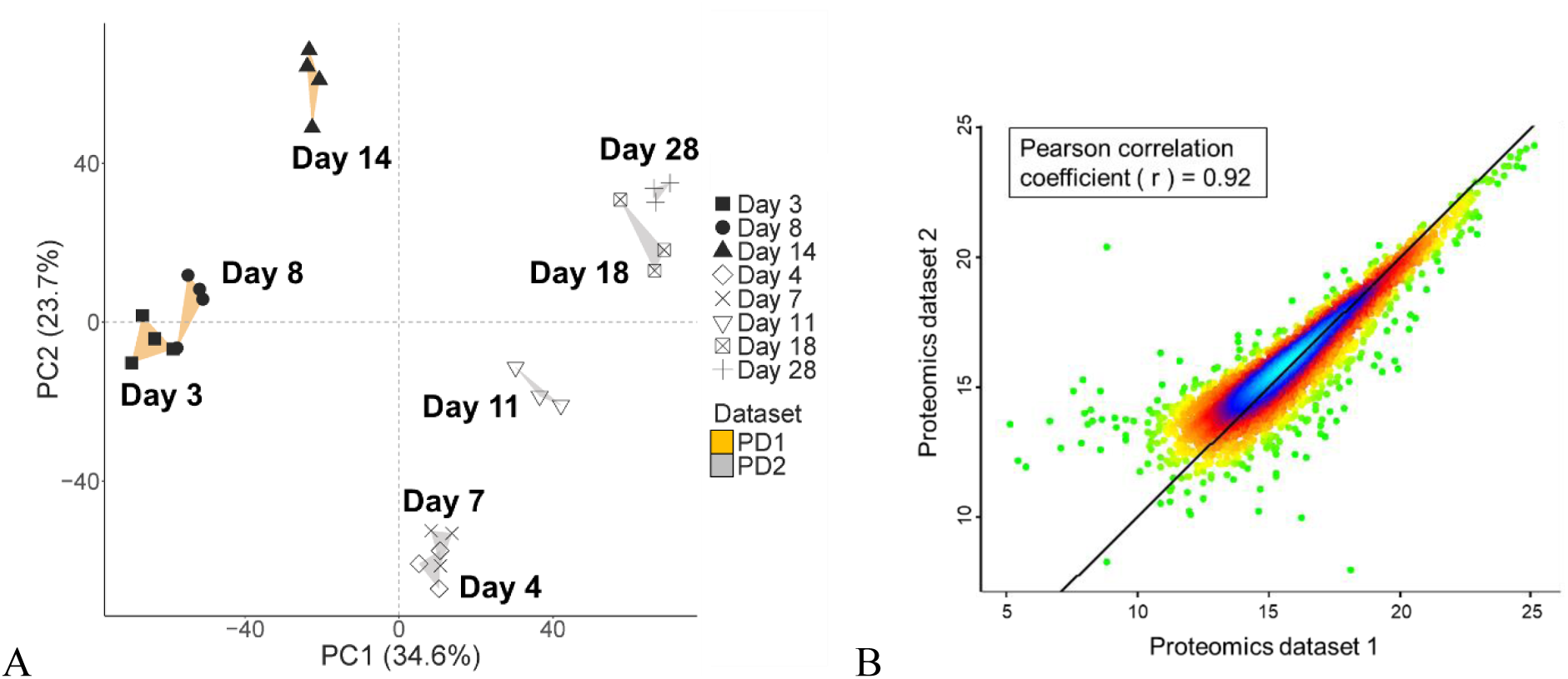
Comparison of protein expression trends in proteomics datasets 1 and 2 (PD1 and PD2). (A) Principal component analysis (PCA) was performed on filtered, log₂-transformed, and zero-centred data, providing 6803 PAC2 protein groups shared between PD1 and PD2. Convex hulls (orange and grey areas for PD1 and PD2, respectively) show the outer boundaries formed by replicates within each group. (B) Correlation and density plot for 6803 PAC2 protein groups shared between PD1 and PD2. For each dataset, abundance values of each protein group were averaged across biological replicates from all conditions (sampling days). The diagonal line indicates equality (x = y); points above the line have higher values in PD2, and those below have higher values in PD1. Color gradient indicates the density of the data points (blue – most dense; red and yellow – intermediate; green – least dense).

The separation of PD1 and PD2 samples could be due to differences in absolute abundance values measured for each protein group in the two datasets, as Pearson correlation coefficient between the two datasets was lower than 1 (r = 0.92, Fig. 3B). These differences appeared to be unevenly distributed within the highest- and lowest-abundance analytes in each dataset, with the former showing slightly higher abundance values in PD1 compared to PD2 and the latter, vice versa, showing slightly lower abundance values in PD1 compared to PD2 (see Fig. S3 for a detailed analysis of top-200 high- and low-abundant protein groups). This uneven distribution of absolute abundance differences could affect the degree of variance within the two datasets, which was likely reflected in the PCA. The differences in the absolute abundance values could be caused by the differences in the peptide amount loaded on the column in chromatographic runs performed for PD1 and PD2 samples (see Table 1), which is reflected by the progressive decrease in Pearson coefficient from 0.964 to 0.827 for comparisons of abundance values between day 3 of PD1 (1 µg loaded) through samples with 2 µg loading (PD1 day 8 and 14) to samples with 4 µg loading (days 4, 7, 11, 18 and 28 of PD2), shown in Fig. S4. For respective comparisons of 2 and 4 µg samples, Pearson coefficients also decreased from 0.939 to 0.841 for day 8 samples, but for day 14 samples, they remained similar or even slightly increased (from 0.86 to 0.886). Of note, correlation between samples is expected to be less than perfect, because they represent different growth phases, accompanied by differential expression of a significant proportion of protein groups (see below). Hence, the observed correlation coefficients can still be considered high overall. It is also important to emphasize that, despite these differences in absolute values, the relative differences between the samples corresponding to exponential and stationary cell population growth phases within each dataset appear to be retained, as shown by similar over-time PCA trends discussed above (Fig. 3A and Fig. S5). Thus, PCA suggests that PD1 and PD2 data are comparable but cannot be directly combined for quantitative analysis aimed at identifying differentially expressed proteins. We therefore proceeded to perform differential expression analysis for PD1 and PD2 separately, followed by comparing the obtained results.

We identified 4450 differentially expressed protein groups (DEPs) in PD1 (corresponding to 62% of quantified proteome, SF 2C) and 3531 DEPs in PD2 (49% of quantified proteome, SF 3C), presented in Fig. 4 as hierarchically clustered heatmaps. A total of 2520 DEPs were shared between the two datasets PD1 and PD2, and they turned out to represent the core of the two main co-expression groups progressing from early to late cell population growth phases, identified as downregulated (1247, Fig. S6A, S6B) and upregulated (1273, Fig. S6C, S6D) DEPs. These shared DEPs were then subjected to functional annotation and enrichment analysis to identify the affected biological processes and functions. To perform the analysis, DEPs were converted to gene symbols, resulting in 1229 genes from 1247 DEPs in the downregulation cluster and 1270 genes from 1273 DEPs in the upregulation cluster. Selected results of functional annotation and enrichment analysis are shown in the top panels (proteomics) of Fig. 5A and 5B, corresponding to the downregulation and upregulation clusters, respectively; the complete outcome of DAVID GO analysis can be viewed in SF 4. Enriched pathways identified in this dataset were related to four main areas: nucleic acid metabolic process and DNA replication; ribosome biogenesis and protein folding; cell cycle process and its regulation; and exocrine system/gland development.

**Figure 4.**
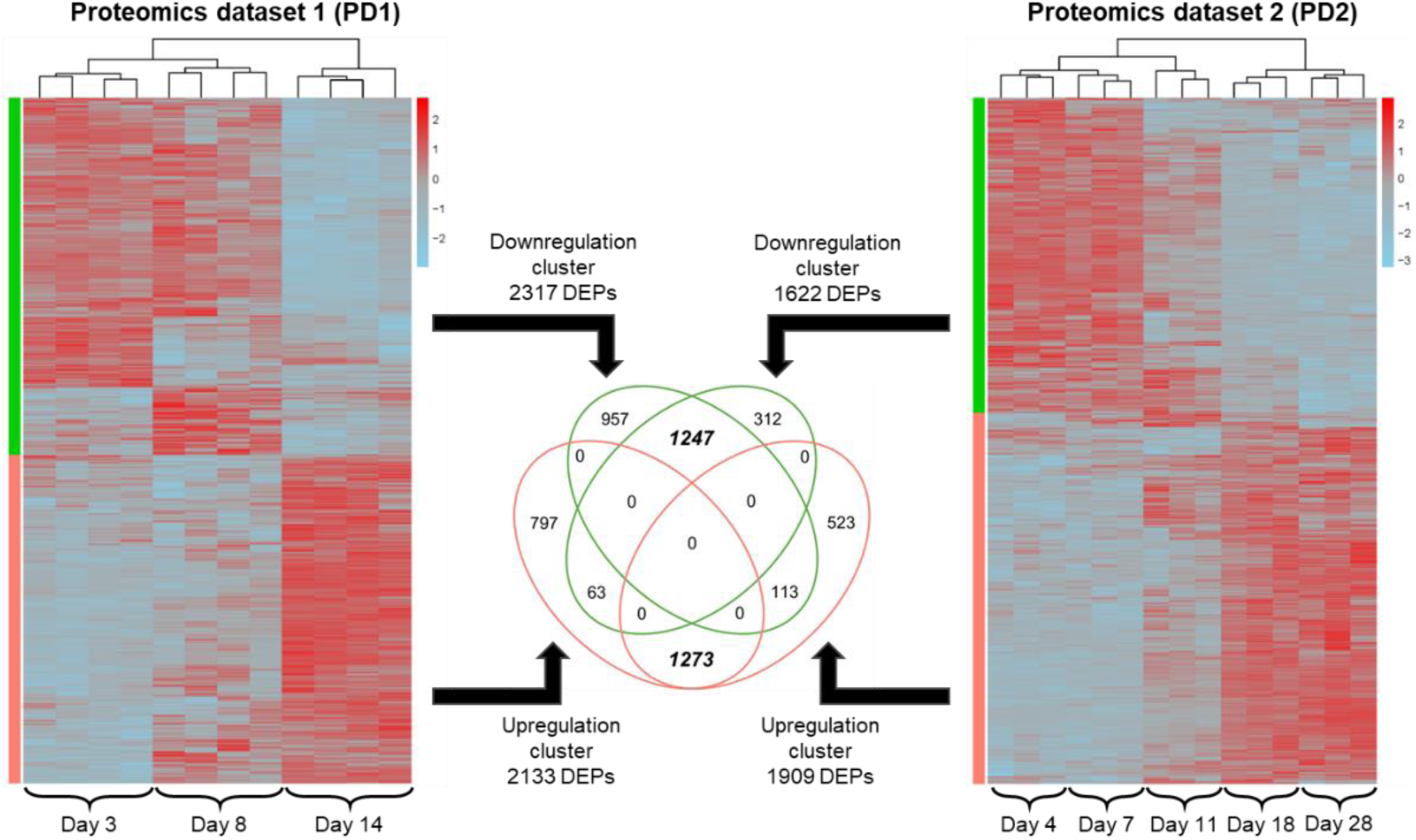
Hierarchical clustering of the differentially expressed protein groups (DEPs) identified in the proteomics dataset 1 (PD1) and 2 (PD2). Color gradient represents the DEPs z-score normalized expression values. Heatmap columns correspond to the biological replicates within each dataset (n=4 for PD1 and n=3 for PD2) and each cell population growth phase (represented by sampling days 3, 8, 14 in PD1 and 4, 7, 11, 18, 28 in PD2), indicated at the bottom of each heatmap.

**Figure 5.**
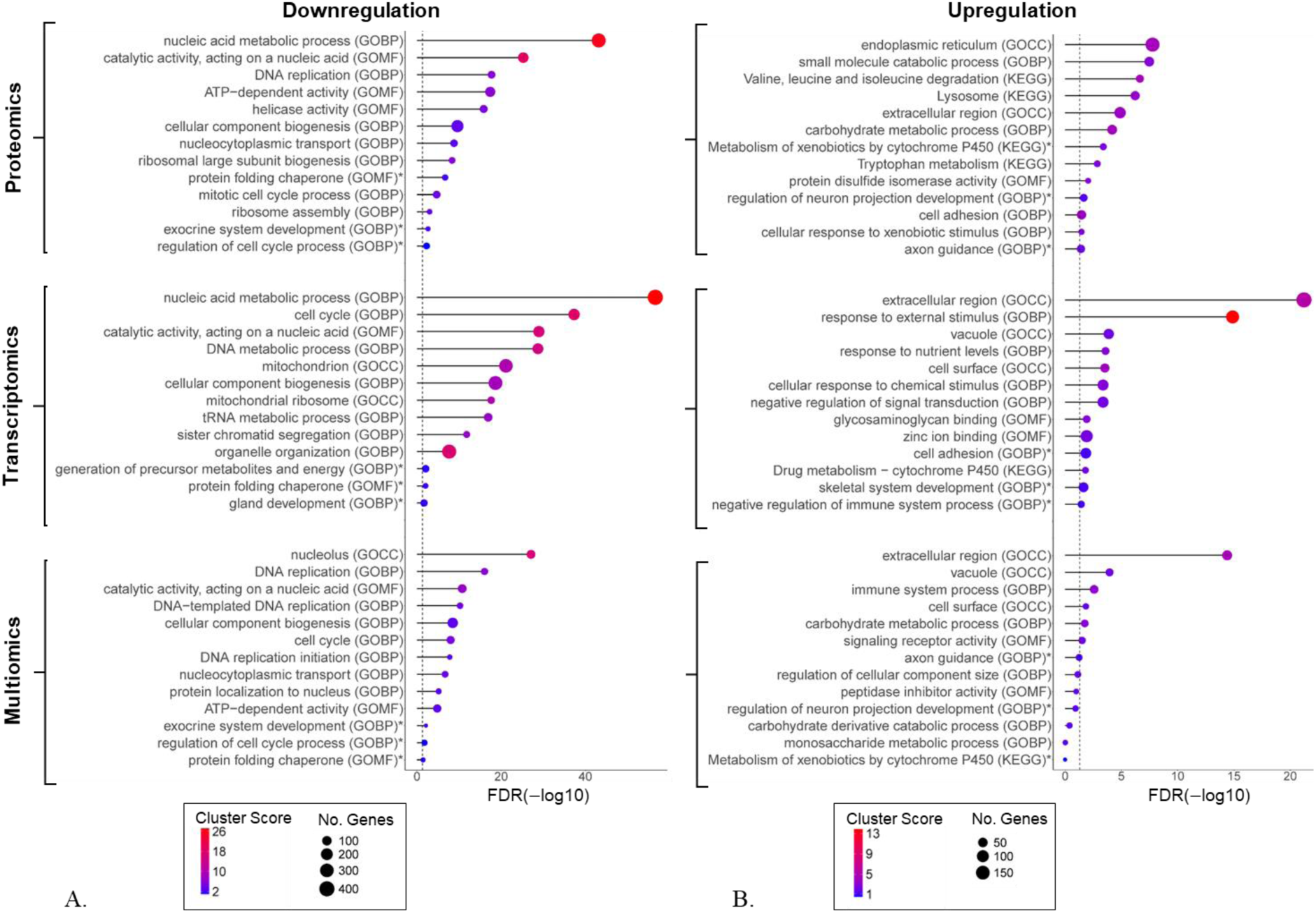
Functional annotation and enrichment analysis of differentially regulated gene clusters across cell population growth phases. Panels (A) and (B) show results for downregulation and upregulation co-expression clusters derived from proteomics analysis (top two graphs, analysis performed on 1247 downregulated and 1273 upregulated differentially expressed proteins (DEPs) shared between proteomics dataset 1 (PD1) and proteomics dataset 2); transcriptomics analysis (middle two graphs, analysis performed on 2306 downregulated and 2143 upregulated differentially expressed genes (DEGs)); and multiomics analysis (bottom two graphs, analysis performed on 690 downregulated and 446 upregulated genes found in both DEPs and DEGs lists from the matched multiomics dataset). Annotation was performed with KEGG pathways and gene ontology (GO) terms for biological process (BP), cellular component (CC) and molecular function (MF). Resulting pathways and terms were analyzed for enrichment against background proteome and/or transcriptome, and clustered based on gene similarity in DAVID Bioinformatics. Shown clusters include top 10 and several additionally selected (indicated by *) statistically significant terms/pathways of interest. The complete outcome of these analyses can be viewed in Supplementary Files (SF) 4, 7 and 8, corresponding to proteomics, transcriptomics and shared omics analysis data. Cluster score represents the geometric mean (-log p-value) of all GO terms and KEGG pathways included within a cluster. Cluster names and gene numbers were taken from the top significant pathway or term (sorted on y-axis based on false discovery rate, FDR, shown on x-axis).

### Transcriptomics profiling of PAC2 cells in exponential and stationary phases

Transcriptomics analysis was performed for matched samples corresponding to day 8 and day 14 within PD1 (see SF 5), representing exponential and stationary cell population growth phases, respectively. RNAseq analysis of PAC2 cell samples generated counts for 30628 gene IDs in total. Following data filtering for very low-expressed genes, the initial transcriptomics dataset (TD) was reduced to 17570 genes (see SF 6A) and finally to 14574 transcripts that remained after filtering with a threshold of transcripts per million (TPM) >1 (see SF 6B). Over 90% (13409) of identified transcripts were shared between the two sampled conditions, i.e., days 8 and 14 (Fig. 6A). These conditions, however, showed a clear separation in the PCA (Fig. 6B), indicating the occurrence of major expression changes, similar to those observed in the proteomics data (see Fig. 3A and Fig. S5).

**Figure 6.**
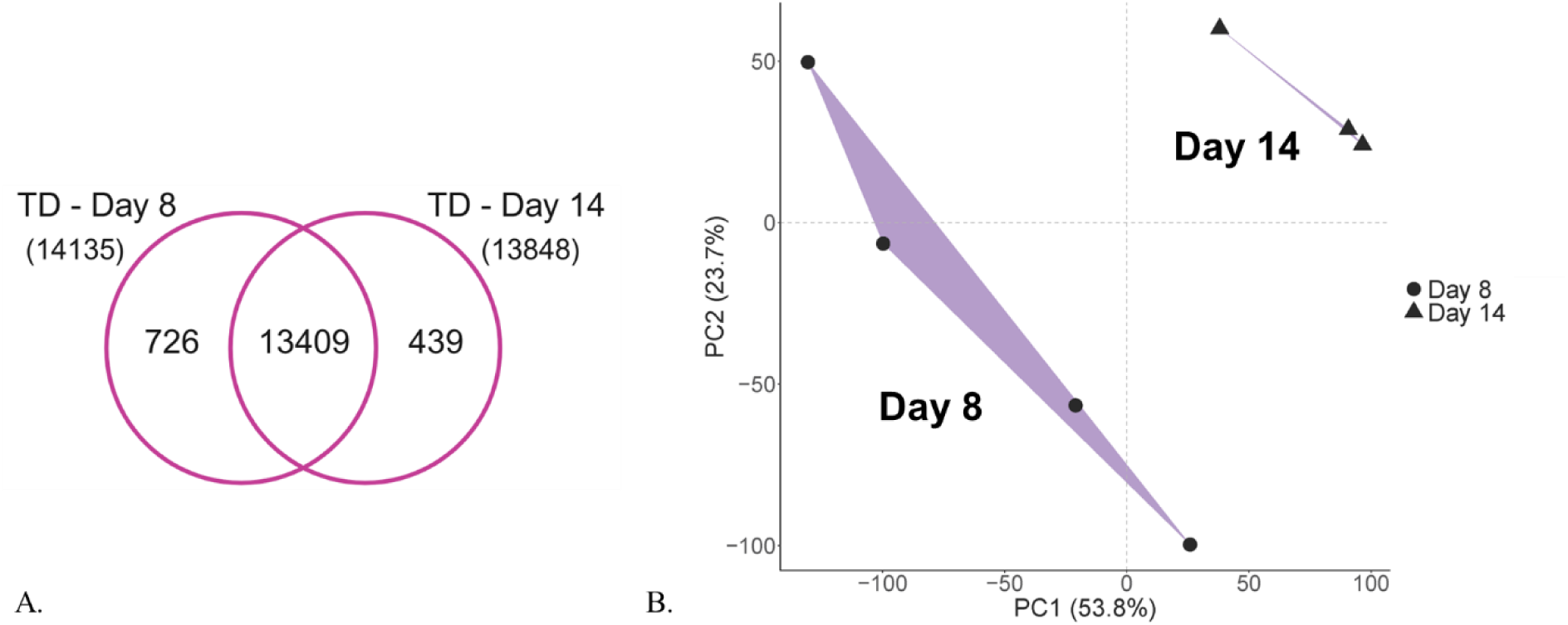
Characterization of the transcriptomics dataset (TD). (A) Numbers and overlap of transcripts identified in the PAC2 cells sampled at two different cell culture growth phases (i.e., day 8 and day 14). In total, 14574 gene transcripts were reliably measured. (B) Principal component analysis (PCA) of the 14574 gene transcripts showed a clear separation between the two different growth phases (day 8 and day 14). Convex hulls (purple areas) show the outer boundaries formed by replicates within each group. Each data point represents a biological replicate. Note that each condition was intended to be analyzed in 4 biological replicates, but in the day 14 condition, one replicate was accidentally lost during library preparation.

Differential expression analysis identified 2306 downregulated and 2143 upregulated differentially expressed genes (DEGs) in the TD (see SF 6C). For the TD cluster of downregulated genes, enriched terms showed a high overlap with all four main areas revealed in the respective proteomics cluster, but there were also several new terms appearing among the top-enriched, including organelle organization, mitochondrion and generation of precursor metabolites and energy (compare top and middle panel in Fig. 5A and see SF 4A and SF 7A). For upregulated genes, PDs and TD clusters share enriched terms related to response to chemical stimulus, cytochrome P450 drug metabolism, extracellular region, cell surface, and adhesion (compare top and middle panel in Fig. 5B and see SF 4B and SF 7B). In contrast, terms related to axon projection development appeared only in upregulated DEPs (Fig. 5B, top panel), while upregulated DEGs showed enrichment of skeletal system development terms and negative regulation of immune system process (Fig. 5B, middle panel).

### Proteomics and transcriptomics provide comparable information about biological changes in PAC2 cells

To explore the overlap between proteomics- and transcriptomics-derived information about molecular and cellular changes occurring upon transition from exponential to stationary cell population growth phase in PAC2 cells, we carried out a differential gene expression analysis of day 8 vs day 14 condition in PD1 and TD and compared the identified DEPs and DEGs. For PD1, this analysis identified 1668 downregulated and 1571 upregulated DEPs (Fig. 7), which shared 98% and 95% overlap with the respective DEP clusters identified in the larger proteomics analysis described in Fig. 4. The terms identified by functional annotation and pathway enrichment analysis of these two clusters were also similar to those highlighted in the complete proteomics data (shown in Fig. 5, top panel).

**Figure 7.**
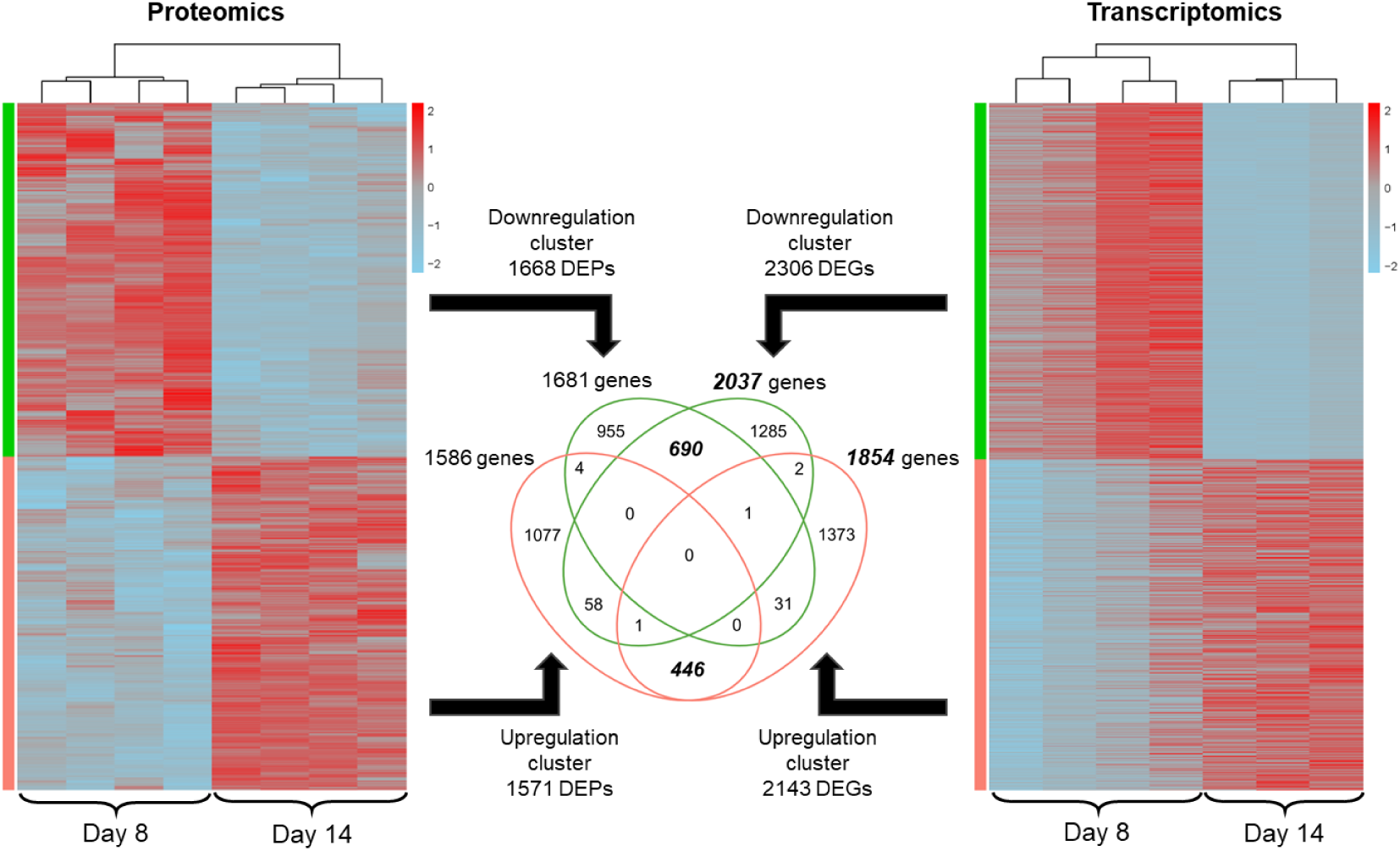
Hierarchical clustering of differentially expressed protein groups (DEPs) and differentially expressed genes (DEGs) in the matched multiomics dataset. Color gradient represents the DEPs’ and DEGs’ z-score normalized expression values. Heatmap columns correspond to the biological replicates (n=4) within each dataset and each of the two analyzed cell population growth phases (i.e., days 8 and 14), indicated at the bottom of each heatmap. Replicate B3 in day 14 condition of TD was not available (accidental loss).

To compare these DEPs with the DEGs identified in the TD (described in the preceding section), both lists were converted from gene identifiers originally used in each dataset (i.e., UniProt accessions in PDs and Ensembl gene IDs in the TD) into official gene symbols. This resulted in slightly different output numbers than were input (compare numbers shown in the middle area of Fig. 7), which could be due to some IDs being less comprehensively annotated, leading to many-to-one or ambiguous mapping. Moreover, a few genes appeared as shared between down- and upregulation clusters within proteomics (4 genes) and transcriptomics (2 genes) lists. This could be due to the ambiguities introduced during the conversion from respective identifiers to gene symbols, but it could also imply the existence of (unresolved) isoforms of the same gene/protein that behave differently.

690 genes were found to be significantly downregulated (FDR<0.05) on both the protein and transcript levels, representing ca. 40% of downregulated DEPs and about one third (ca. 34%) of downregulated DEGs (Fig. 7). Overlap between proteomics and transcriptomics data was even lower within the upregulated cluster, with 446 shared genes representing ca. 28% and 24% of all upregulated DEPs and DEGs, respectively (Fig. 7). However, despite this low overlap and the relatively low numbers of shared genes, the terms enriched among the shared DEPs and DEGs in the downregulation cluster (see Fig. 5A, bottom panel, and SF8A for complete analysis output) were found to cover the same four biological groups that were identified as downregulated based on the individual analyses of proteomics (Fig. 5A, top panel) and transcriptomics (Fig. 5A, middle panel) data, i.e., DNA replication, translation, metabolism and cell cycle regulation. Similarly, enriched terms in the shared upregulated DEPs and DEGs included terms related to extracellular region, response to chemical stimulus and drug metabolism (Fig. 5B, bottom panel), also identified in the individual proteomics (Fig. 5B, top panel) and transcriptomics (Fig. 5B, middle panel) analysis. Interestingly, the terms enriched in the shared proteomics/transcriptomics list also included both the axon guidance and immune system process - the two areas found to be uniquely enriched in the individual proteomics and transcriptomics lists, respectively (compare bottom, top and middle panels in Fig. 5B), when analysed separately. The complete output of functional annotation and enrichment analysis for shared DEPs and DEGs can be viewed in SF 8A and SF 8B for downregulated and upregulated clusters, respectively.

Top-changing genes (identified as showing the log₂ difference >|1.5| and p-value <0.01, see Fig. 8) included 60 DEPs and 136 DEGs downregulated and 51 DEPs and 343 DEGs upregulated (see SF 9A and SF 9B for complete lists of DEPs and DEGs, respectively). Among these, 5 downregulated and 16 upregulated genes were shared between DEP and DEG lists (Fig. 8 and Table 2).

**Figure 8.**
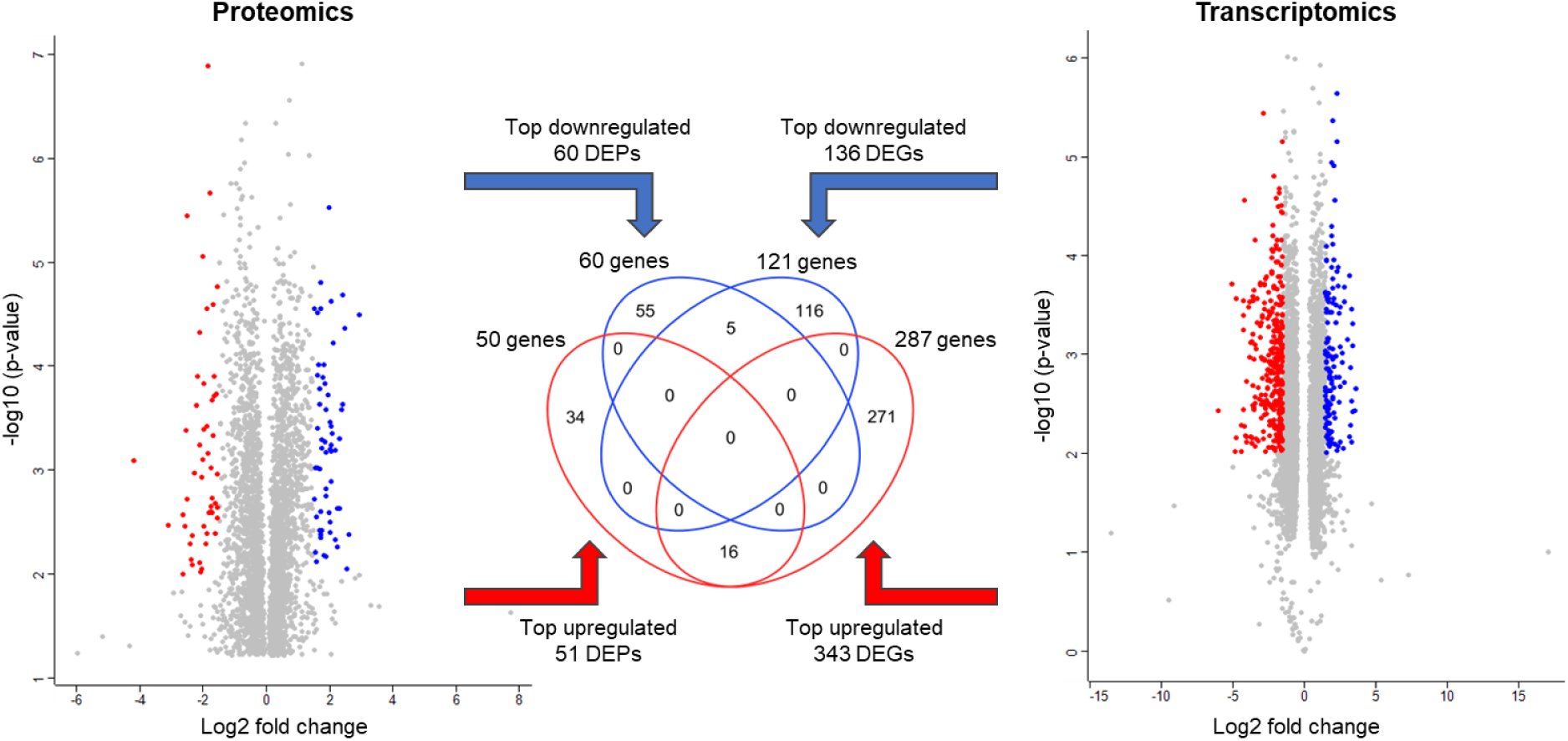
Overlap between the top regulated differentially expressed protein groups (DEPs) and transcripts (DEGs) in PAC2 cells from day 8 to day 14. The lists of 3239 DEPs from proteomics dataset 1 (PD1, data from days 8 and 14) and 4449 DEGs from the transcriptomics dataset (TG) were tested using Student’s t-test and plotted in volcano plots shown in left and right panel, respectively. Identifiers with increased expression (log₂ difference >1.5 and p<0.01) are shown in red, corresponding to 51 DEPs (A) and 343 DEGs (B). Identifiers with decreased expression (log₂ difference <–1.5 and p<0.01) are shown in blue, corresponding to 60 DEPs (A) and 136 DEGs (B). Grey dots represent the remaining DEPs and DEGs not meeting these thresholds. To identify overlaps, protein group identifiers (UniProt accessions) and transcript identifiers (Ensembl IDs) were mapped to gene symbols using DAVID Bioinformatics.

**Table 2.**
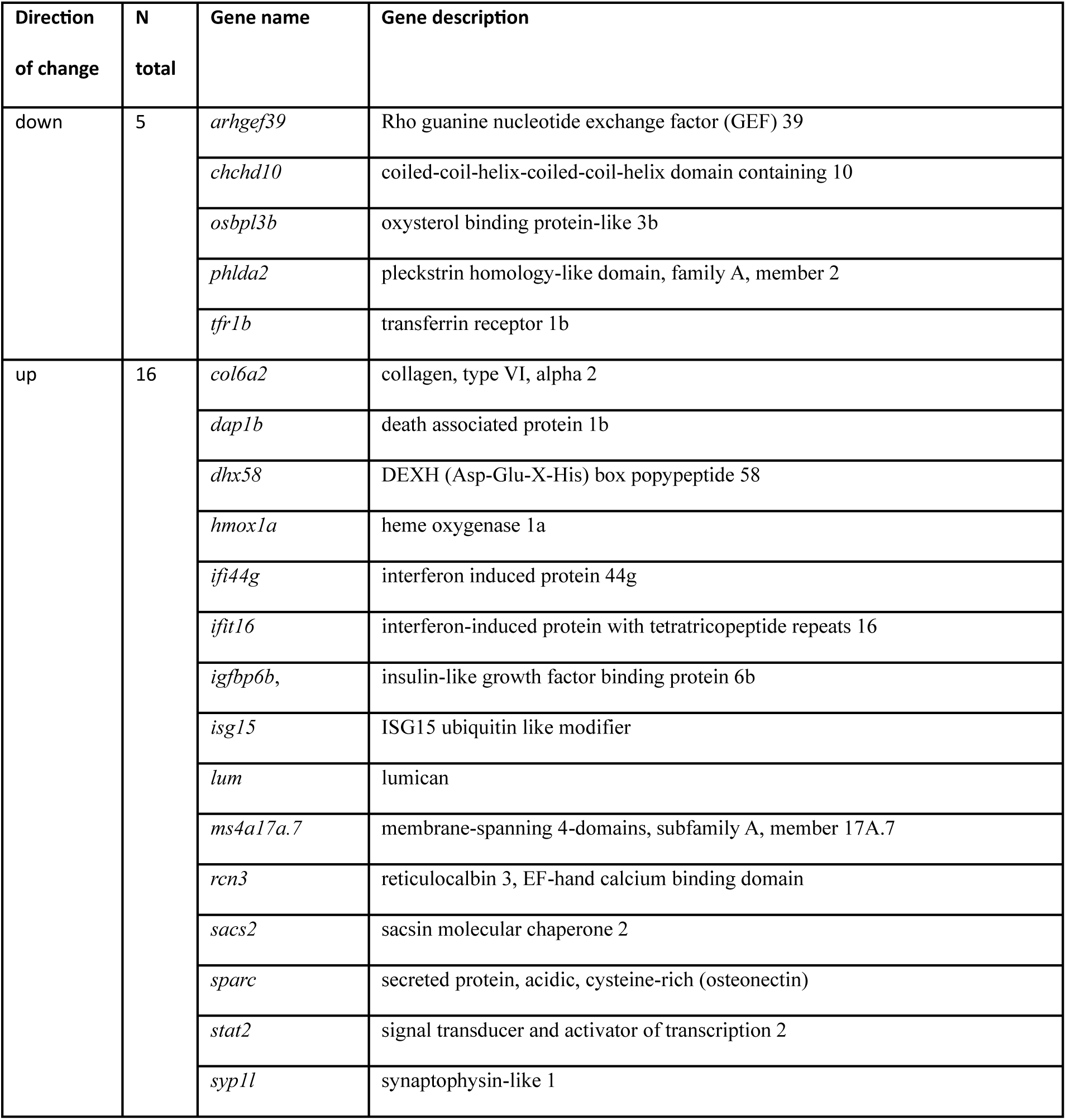
Shared genes found on both the protein and transcript lists of top-regulated exponential-to-stationary genes. Significantly regulated genes were identified by Student’s t-test and top-regulated genes were identified as showing the log₂ difference >|1.5| and p<0.01. Overlaps between differentially expressed proteins and gene transcripts were identified by comparing respective lists converted to official gene symbols.

### Multiomics integration model of gene expression changes between exponential and stationary phases

An unsupervised multiomics model, built to evaluate the shared variance between the proteomics and transcriptomics datasets, showed a clear separation between day 8 and 14 (Fig. 9A). This trend was similar to that observed in proteomics (Fig. 3A) and transcriptomics (Fig. 6B) data individually. In the model, each biological replicate exhibited relatively small variance differences between the proteomics and the transcriptomics dataset, with the highest differences observed in biological replicate 4 (Fig. 9A). Contribution to the total explained variance per component was nearly equal for transcriptomics and proteomics (Fig. 9B).

**Figure 9.**
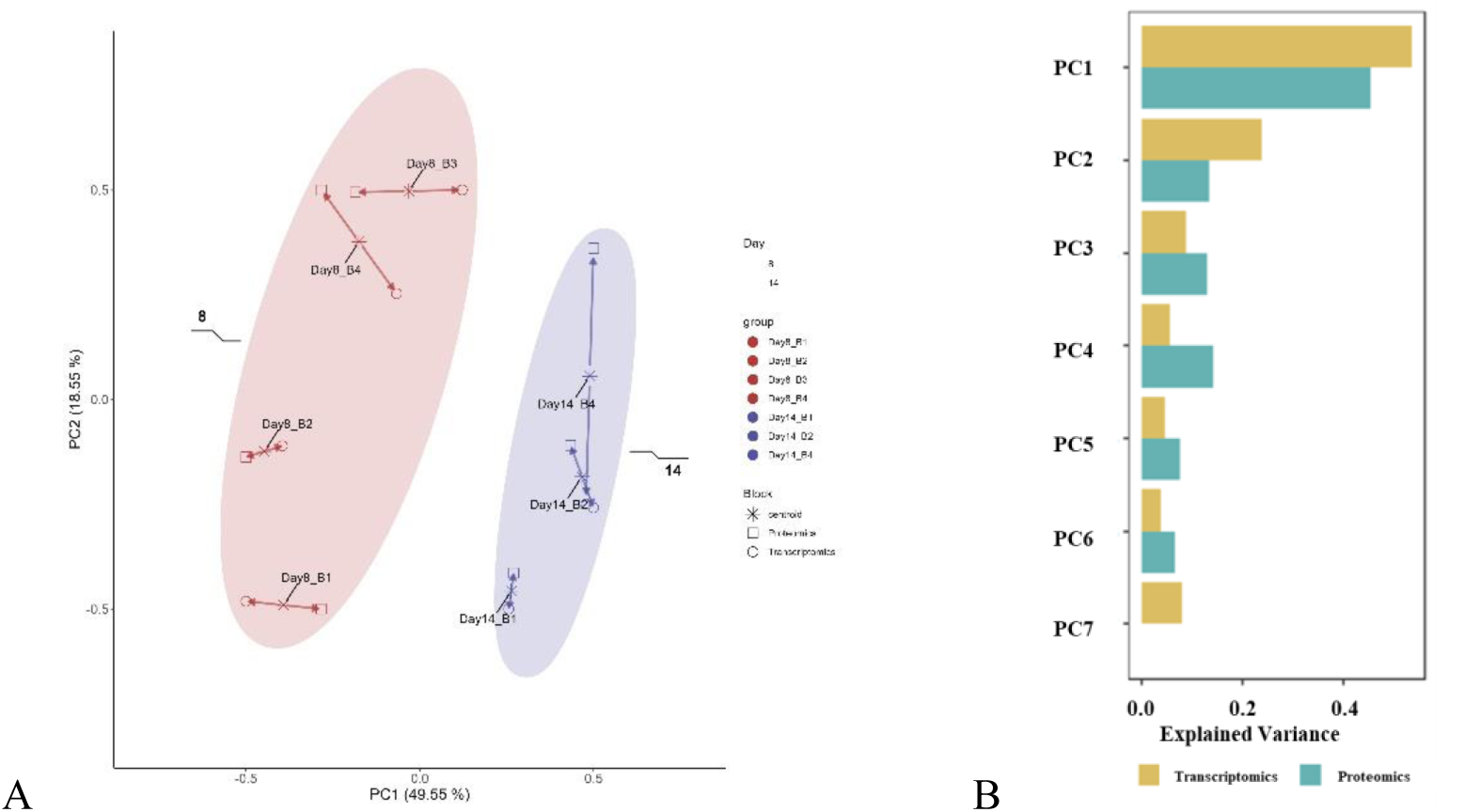
Multiomics Partial Least Square (PLS) model showing separation between analyzed conditions corresponding to exponential and stationary cell population growth phases. Multiomics model was built for data from days 8 and 14 from proteomics dataset 1 (PD1) and the corresponding matched samples from the transcriptomics dataset (TD). Arrows represent the variance difference between proteomics and transcriptomics datasets for each biological replicate, with shorter arrows indicating stronger agreement in variance. Panel (B) shows contributions of proteomics and transcriptomics data to the total explained variance per principal component (PC).

### Functional annotation analysis of all proteins and transcripts detected in PAC2 cells

Following the above-described analysis of DEPs and DEGs that accompany the transition from exponential to stationary growth phase in PAC2 cells, we wanted to explore which types of biological processes and functions, especially the toxicologically relevant ones, are being expressed in PAC2 cells overall. For this we carried out a functional annotation analysis of all proteins and transcripts detected in PAC2 cells. To enable comparisons, original identifiers were converted to official gene symbol identifiers. As already observed previously, the output of this conversion did not yield the exact number of identifiers that were input. That is, 7320 protein groups (PD1 and PD2 combined) resulted in 7465 genes, while 14574 transcripts resulted in 12172 genes. 6644 genes were shared between the complete proteomics (combined PD1 and PD2) and transcriptomics (TD) identifications, corresponding to 89% and 55% of PDs and TD. Thus, around 11% (821) of protein groups detected by global proteomics did not have a corresponding transcript match in the RNAseq data. Functional annotation analysis of these genes in DAVID GO identified 6 clusters with significantly enriched terms related to extracellular region and immune response; cell adhesion; ligand-gated channel activity and neurotransmitter receptor activity; DNA-binding transcription factor activity; supramolecular fiber; and calcium signaling (see SF 10). Conversely, over half of the identified transcripts were also detected on the protein level, while around 45% were identified on the transcript level only, which is to be expected in multiomics studies since transcriptomics typically shows higher detection sensitivity than proteomics.

Functional annotation analysis of all proteins (SF11 A) and transcripts (SF11 B) detected in PAC2 cells showed that, despite the almost 2-times difference between the overall gene counts present in the proteomics and transcriptomics datasets, similar functional categories were identified by both data types (compare Fig. 10A and Fig. 10B). Additional information about each functional category is provided in Table 3, and a complete overview of Cytoscape categories can be seen in SF 11C.

**Figure 10.**
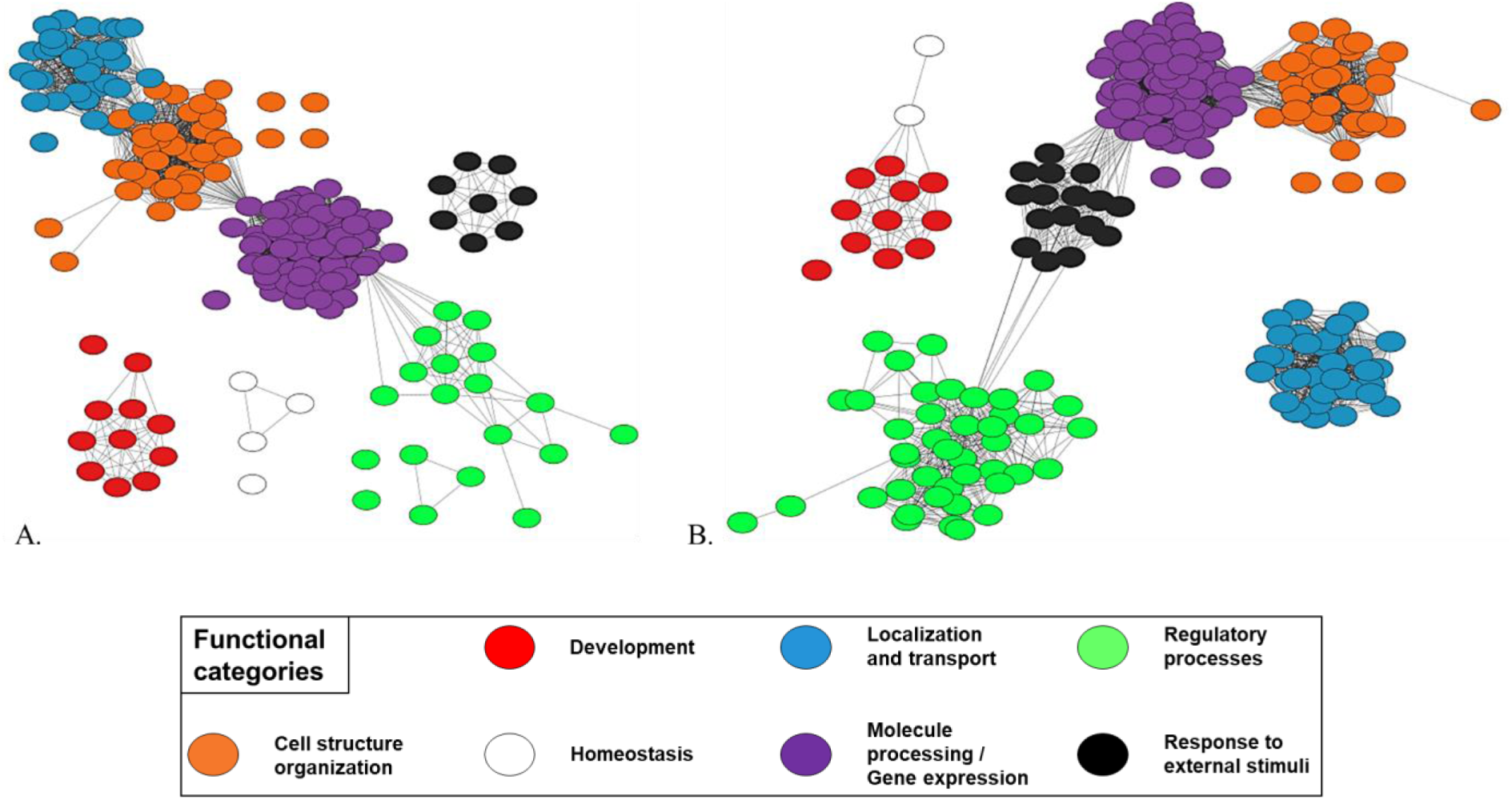
Overview of the main biological processes represented by the proteins and transcripts measured in zebrafish PAC2 cells. Panel (A) shows representative gene ontology biological process terms identified within the 7465 genes corresponding to the 7320 protein groups of the PAC2 proteome. Panel (B) shows the same for the 12172 genes corresponding to the 14574 transcripts of the PAC2 transcriptome. Nodes represent GO biological process terms, and edges connecting them are based on semantic similarity scores calculated in Revigo. Only terms with similarity above the set threshold (0.7) are connected. Unconnected nodes were retained because they were considered informative, despite low similarity to other terms. Colors used to denote different functional categories explained in the shared legend underneath panels (A) and (B) were manually assigned based on proximity and nomenclature (shared common words).

**Table 3.** Overview of the main functional categories containing enriched terms and pathways represented in the proteome and transcriptome of PAC2 cells. For each category, the number of terms, the top statistically significant term (marked by *) and the most mapped term with its corresponding number of genes are shown.

| Functional categories | Proteome-based | Transcriptome-based |
| --- | --- | --- |
| **Cell structure organization** | - 42 terms<br>- ribonucleoprotein complex biogenesis*<br>- organelle organization (960 genes) | - 37 terms<br>- ribonucleoprotein complex biogenesis*<br>- organelle organization (1449 genes) |
| **Development** | - 11 terms<br>- chordate embryonic development*<br>- embryo development (477 genes) | - 11 terms<br>- convergent extension*<br>- embryo development (718) |
| **Homeostasis** | - 4 terms<br>- cellular homeostasis*<br>- homeostatic process (292 genes) | - 2 terms<br>- multicellular organismal-level homeostasis (181 genes)* |
| **Localization and transport** | - 34 terms<br>- cellular localization (858 genes)* | - 35 terms<br>- cellular localization (1158 genes)* |
| **Molecule processing /**<br>**Gene expression** | - 80 terms<br>- macromolecule biosynthetic process*<br>- protein metabolic process (1391 genes) | - 82 terms<br>- nucleic acid biosynthetic process*<br>- protein metabolic process (2160 genes) |
| **Regulatory processes** | - 19 terms<br>- regulation of translation*<br>- regulation of cellular component organization (363 genes) | - 42 terms<br>- regulation of catabolic process*<br>- regulation of DNA-templated transcription (1358 genes) |
| **Response to external stimuli** | - 8 terms<br>- cellular response to stress (387 genes)* | - 16 terms<br>- cellular response to stress*<br>- response to stress (1022 genes) |

### PAC2 cell type and relevance of PAC2 cells for toxicological studies, explored by multiomics analysis

Manual checks of selected gene families of interest were carried out to gain information on the PAC2 cell type as well as to explore the relevance of PAC2 cells for toxicological studies. First, we looked at the expression of selected markers of fibroblastic or epithelial cell types, specifically the fibroblastic markers: *vim*, vimentin; *cdh2*, cadherin 2, type 1, N-cadherin (neuronal); *col1a1b,* collagen, type I; *fn1a*, fibronectin; and epithelial markers: *cdh1*, cadherin 1, type 1, E-cadherin (epithelial); *prr36b*, mucin- 5AC; *si:ch73-61d6.3*, occludin; *sdc2*, syndecan; *tjp2b*, zona occludens 2. All markers except vimentin were found to be expressed in PAC2 cells at both the protein and transcript level (Table 4 and Fig. S7). Next, we looked at the expression of selected toxicologically relevant gene families and pathways, also summarized in Table 4. PAC2 cells were found to express molecular components related to cytoprotective mechanisms (e.g., detoxification and response to heat shock response, oxidative stress, endoplasmic reticulum stress), which are often involved in responses to toxic chemicals (36, 37). With regard to xenobiotic metabolism, PAC2 cells express several members of gene families related to active transport (ABC, SLC) and biotransformation, including phase I: cytochrome P450 (CYP); phase II: glutathione S-transferases (GST); uridine 5-diphospho-glucuronosyltransferases (UGT), sulfotransferases (SULT); and N-acetyltransferases (NAT). Further, PAC2 cells also express a number of toxicologically relevant signaling pathways, such as MAPK/JNK, AKT and mTOR, as well as transcription factors and effector genes involved in response to hypoxia and metals, heat shock response, unfolded protein response and autophagy regulation. Lastly, PAC2 cells express a number of nuclear receptors, including those that govern the response to important chemical groups, including estrogens, glucocorticoids, aryl hydrocarbons, and peroxisome proliferator substances. While expression was most often observed on transcript level only, numerous genes were detected on both the transcript and protein level, and several more were detected on protein level only (Table 4).

**Table 4.**
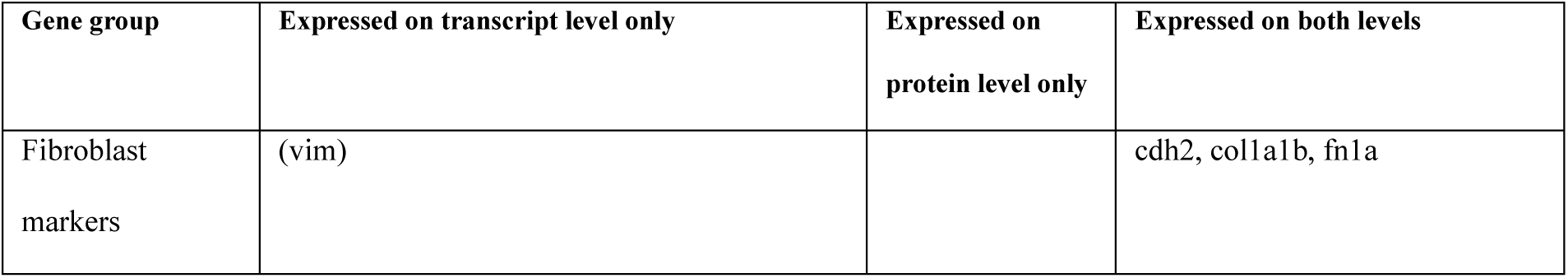
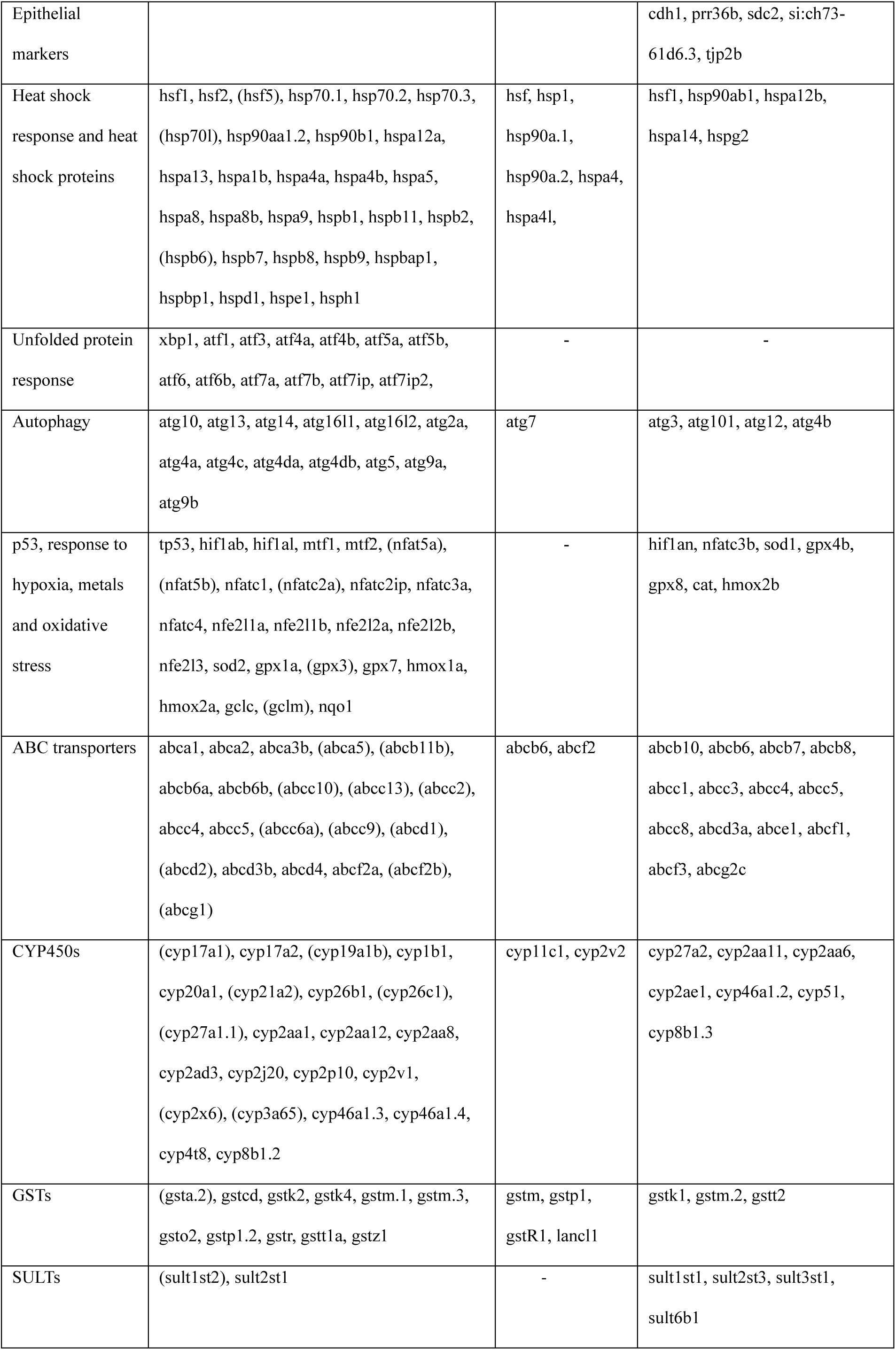
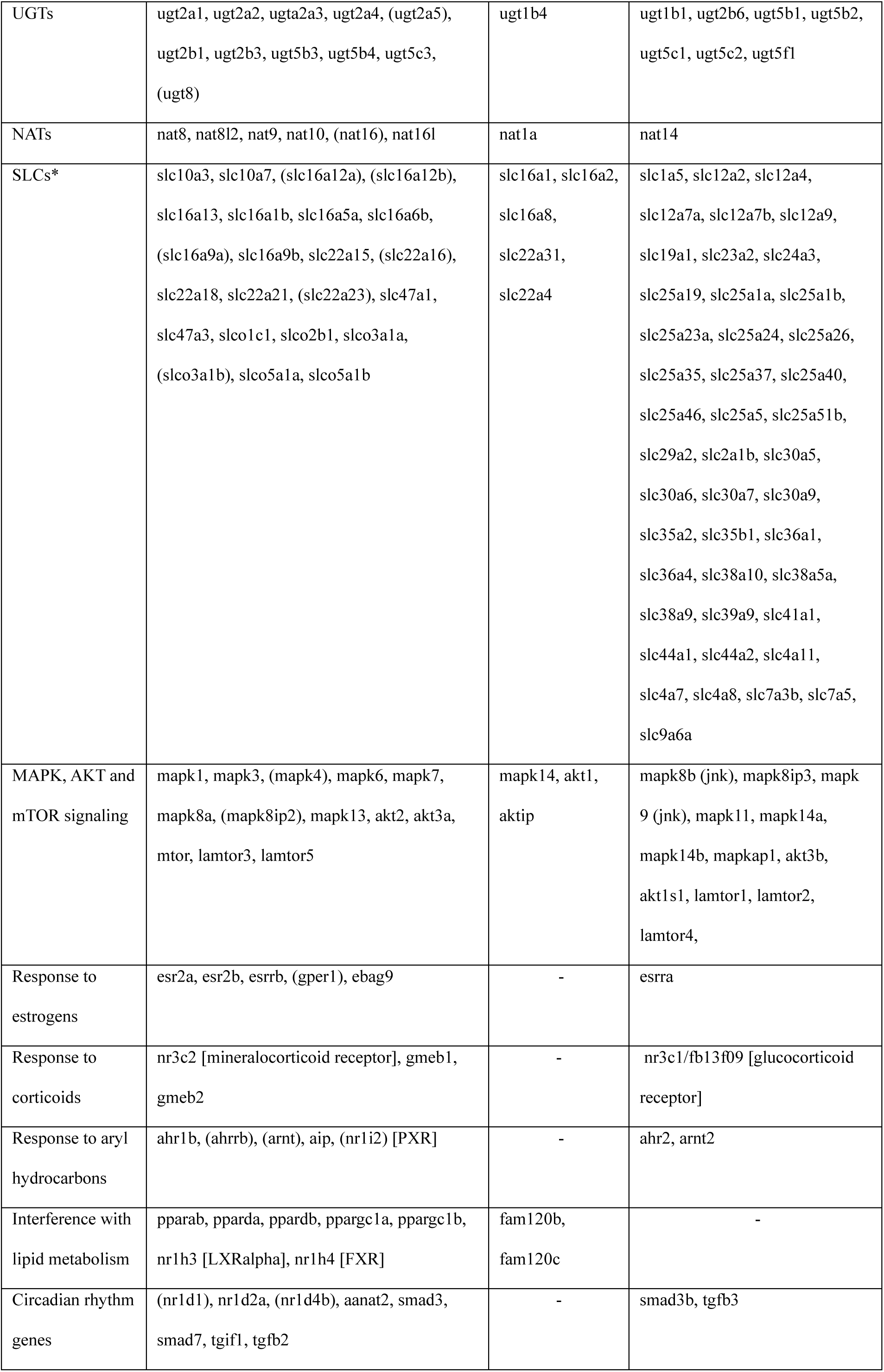
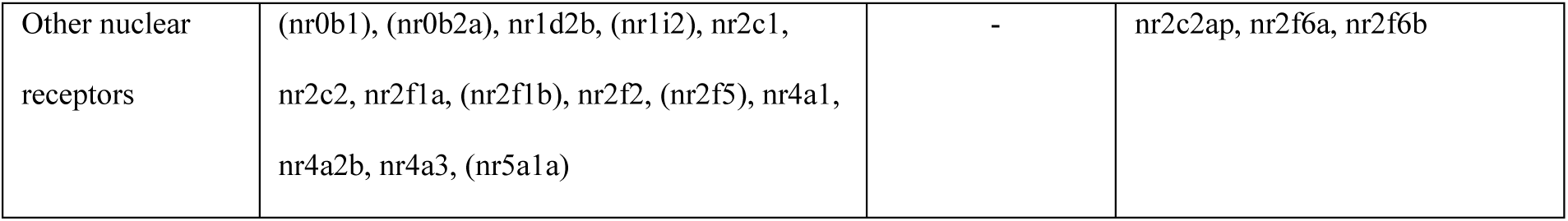
Overview of expression in PAC2 cells of genes or proteins belonging to selected families related to toxicologically relevant processes and functions. Selected gene families were checked by manual search in proteomics datasets (PDs) and transcriptomics dataset (TD) for respective keywords or gene symbols. *For SLC, only selected families known to be involved in transport of xenobiotics plus genes whose expression was detected on both the mRNA and protein level, are shown. Gene names shown in brackets () indicate genes that were detected only in the TD’s counts list. Receptor names are indicated in square brackets [] if the identity if not obvious from gene name.

## Discussion

We carried out a comprehensive profiling of molecular changes occurring in PAC2 cells during progression through the cell population growth phases. Extensive data obtained on the protein level using two distinct sets of samples allowed evaluating the robustness of global proteomics analysis. Proteomics analysis (PD1, PD2) was complemented with transcriptomics data generated at two time points in exponential and stationary growth phases (TD), which were directly paired with corresponding samples from PD1. The inclusion of transcriptomics analysis allowed increasing the depth of gene expression coverage, which was important for functional characterization of the cells. It also permitted to compare the overall molecular insights provided by the two omics data streams with regard to gene expression changes occurring between growth phases and supported the in-depth evaluation of the overall repertoire of genes and pathways expressed in the PAC2 cells.

### Data independent acquisition (DIA)-based global proteomics enables fast analysis and deep coverage

Total number of PAC2 protein groups measured in our study was 7320, with over 95% overlap observed across all technical and biological replicates. This comparatively deep coverage was achieved by separating the complete peptide sample (i.e., without prior fractionation) with a ca. 2h nanoLC gradient on in-house made 30 cm-long column, coupled to MS/MS in data independent acquisition (DIA) mode (38), followed by directDIA (library-free) analysis (39) to identify and quantify peptides and protein groups using Spectronaut^TM^ software (40, 41). The speed and depth of proteome analysis achieved here constitute a significant improvement compared to previously published studies on proteomes of cell lines from zebrafish and other fish species, which typically reported 2-5 times lower coverage for unfractionated samples (42–44), while a more comparable coverage has been achieved by including extensive fractionation steps, e.g., in a study with early zebrafish embryos (45). The improvement reported here is largely based on the implementation of DIA mode, which offers superior sensitivity and more stable label-free quantification results compared to the data dependent acquisition (DDA) more commonly used in earlier studies (38, 46, 47). The fast and sensitive global proteomics analysis workflow presented in this study paves the way for a broader application of global proteomics in ecotoxicological studies, especially those that require analyzing large sample numbers to obtain time- and concentration-resolved molecular profiles. This is often necessary to gain an improved understanding of toxicity mechanisms for unknown or insufficiently characterized substances. While further improvements in speed and sensitivity can be achieved with transition to the next generation of high-resolution mass spectrometers, such as Thermo Orbitrap Astral (48), the presented solution is already capable of obtaining highly informative proteomics profiles (see below) using more regular high resolution mass spectrometry instrumentation.

### DIA proteomics enables robust profiling of proteome changes across PAC2 population growth phases

Analysis of protein expression changes across PAC2 cell population growth phases was based on two different experiments: PD1, performed with four biological replicates in the current study and partially matched with transcriptomics analysis; and PD2, performed with three biological replicates in the frame of a previously published targeted phosphoproteomics study (35). While both experiments had the same overall goal of collecting PAC2 cell samples at different phases of cell population growth, they had several differences in cell culturing conditions, sample collection days, proteomics sample preparation techniques and the total peptide amount loaded on-column for LC-MS/MS analysis (Table 1). Regarding sample preparation, while the PD1 samples were subjected to the regular S-Trap assisted tryptic digest followed by desalting (49), PD2 samples were additionally subjected to Fe^3+^-NTA phosphopeptide enrichment procedure and represented the non-bound fractions collected during its performance (35, 50). Despite these differences, we observed ca. 98% overlap of protein groups (7165) identified within each dataset, while peptides and precursors showed ca. 95% overlap. Comparable overlaps were also observed across the different cell population growth phases (Figure 2). These observations underscore the high consistency and robustness of global proteomics analysis applied in our study. This also confirms that the two different sample preparation techniques delivered largely comparable peptide populations enabling the identification of the same proteins. Thus, passing the non-phosphorylated peptides through the Fe^3+^-NTA enrichment procedure and recollecting them as the non-bound fraction does not seem to lead to substantial losses that would have precluded identification of corresponding proteins.

Despite the high overlap in total identifications, PCA revealed significant separation between the two datasets, particularly in principal component 1 (PC1, Figure 3A). This separation could be due to dataset-specific differences in the absolute expression values, observed especially for the proteins located at the extremes of the dynamic abundance range, i.e., either very high- or very-low abundant proteins (see Fig. 3B and Fig. S3). These deviations could be caused either by variations in peptide recovery introduced during the sample preparation procedure or from different storage regimes or because of the differences in the absolute peptide amounts injected on-column during LC-MS/MS runs (Table 1). Indeed, the coefficient of correlation between different samples showed a tendency to decrease from 0.964 (for 1 µg vs 2 µg loading) to 0.899 (for 1 µg vs. 4 µg loading) in otherwise comparable sample pairs (Fig. S4). However, it also has to be kept in mind that the correlation between samples cannot be expected to reach 100% because each sample is from a different growth phase where some differential expression is naturally present. Therefore, coefficients close to or above 0.9 can still be seen as high, and hence the variations caused by different loading amounts can be considered rather minor, which is also underscored by comparable peptide and protein identifications obtained across all samples (CV 2-5.5%, SF 1).

Lastly, it is important to emphasize that, despite these differences in the absolute abundance values, the two datasets PD1 and PD2 revealed highly similar trends in the temporal protein expression and changes occurring across cell population growth cycle, demonstrated by both the PCA (Fig. 3A) and the comparative analysis of DEPs identified within each dataset, with high overlap between the two datasets observed in the down- and upregulation clusters (Fig. 4, Fig. 5). This again demonstrates that DIA-based global proteomics analysis performed in this study is sufficiently robust against minor differences in cell culture parameters such as plate format, differing sample collection timepoints, or even differences in sample preparation parameters. Moreover, our analysis showed that non-bound fraction from the Fe-NTA phosphopeptide enrichment procedure can be used to obtain reliable global proteomics data. This approach can be applied for studies interested in simultaneous characterization of proteome and phosphoproteome, as it would allow for these two omics streams to be obtained from fully paired samples (essentially, one sample), thus increasing the power for matched multiomics analysis.

### Major gene expression shifts occur upon PAC2 transition from exponential to stationary growth phase

The transition from exponential to stationary cell population growth phase is known to represent a coordinated cellular reprogramming event characterized by major modulation of core metabolic and regulatory networks, reported in both prokaryotes (51) and eukaryotes (52, 53). In PAC2 as well, differential expression analysis revealed massive shifts in both the protein (Fig. 4) and transcript (Fig. 7) expression, indicating comprehensive remodeling of molecular architecture of PAC2 cells across cell population growth phases. The exact timing of exponential-to-stationary transition can be explored based on the characteristics that the cells exhibit over time in culture. The PAC2 growth curves defined either by direct cell count (35) or by Hoechst nuclear staining-based count or by metabolic activity (this study, Fig. 1), suggested that the switch between these phases is likely to occur between day 11 and day 14 post-seeding. By this time, cells are approaching confluence, which is likely associated with a decline in nutrient availability from the culture medium and an increase in cell-to-cell contact interactions. This in turn could trigger a starvation response as well as specific differentiation reactions, which could also accompany the transition from active proliferation to stationary maintenance mode.

These observations are corroborated by the observed changes in the molecular profiles of PAC2 cells across cell population growth phases, revealed in the heatmaps for identified DEPs (Fig. 4, 7) and DEGs (Fig. 7). While coordinated gene expression differences between days 8 (exponential) and 14 (stationary) are clearly visible on both the transcript and protein levels (Fig. 7), a more resolved view can be obtained from the proteomics data shown in Fig. 4 for PD1 and PD2. Here, protein expression patterns at days 3 and 8 (PD1) or 4 and 7 (PD2), corresponding to exponential growth phase, showed clear differences to the patterns observed at day 14 (PD1) and days 18 and 28 (PD2), corresponding to stationary phase. In contrast, day 11, the timepoint present only in PD2, showed an intermediary pattern that significantly differed from both the former and the latter group (but was closer to the former). This suggests that day 11 likely corresponds to a transitionary state between the exponential and stationary phases. Thus, gene expression profiles measured on this day are likely reflective of a switch that the proteome undergoes upon progressing from exponential to stationary phase.

Functional annotation analysis of shared DEPs and DEGs identified between the exponential and stationary growth phases delivered largely similar conclusions with regard to the accompanying changes in the internal molecular machinery of PAC2 cells. That is, both omics types revealed a similar set of affected functions (Fig. 5, top and middle panel), which were also enriched in the core group of shared DEPs and DEGs (Fig. 5, bottom panel). Genes that were downregulated upon transition from exponential to stationary phase covered cellular processes related to DNA replication and nucleic acid metabolic process; mitotic cell cycle process and its regulation; and biogenesis and assembly of ribosome and protein folding chaperones. This suggests a reduction in cell division/proliferation as well as stalled production of cellular structures and organelles, accompanied by reduced translation. The same mechanisms are highlighted by the top-5 genes identified as downregulated with p<0.01 on both the protein and transcript level (Fig. 8). For example, *arhgef39* encodes a RhoGEF protein that regulates cytoskeletal dynamics, cell migration, and proliferation. While previous studies mostly looked at the role of this gene in cancer progression, its downregulation in fish cells may indicate reduced cytoskeletal remodeling, consistent with contact inhibition and decreased proliferation in stationary phase (54, 55). Another downregulated gene, *phlda2*, is a modulator of growth and apoptosis, known in mammals as an imprinted gene involved in growth regulation (56, 57). The *osbpl3b* gene belongs to the family of lipid transfer proteins highly conserved between zebrafish and humans (58). Given the likely role in intracellular lipid trafficking, its downregulation may reflect a decreasing need for membrane turnover, in line with reduced cell growth and proliferation. Similarly, downregulation of the iron uptake receptor *tfr1b* suggests diminishing demand for iron, possibly due to lowered biosynthetic activity typical of quiescent cells (59). In line with this, downregulation of *chchd10* known to be involved in mitochondrial cristae maintenance and regulation of lipolysis, suggests a reduction in energy production, which could reflect a reduced energy demand associated with a metabolic shift toward maintenance rather than growth (60–62).

Processes enriched in the upregulation clusters indicated that cells in the stationary phase prioritize functions related to cellular response to exogenous stimulus and cytoprotective mechanisms, such as detoxification (metabolism of xenobiotics by cytochrome P450) and increased lysosomal activity (autophagy), as well as organelle stabilization and adaptation to the extracellular environment, including extracellular matrix remodeling and changes in cellular adhesion. In addition, stationary-phase cells showed a massive upregulation of immune system-related responses, which will be discussed in more detail in the next section. The transition to the stationary phase can thus be seen as a cellular response to the changing environment, characterized by increased stress due to higher cell density, ensuing nutrient limitation and increased concentration of waste products of metabolism. Thus, robust responses observed at this stage can be indicative of the overall cellular capacity to mount efficient stress response.

Similarly, to the top downregulated genes, most of the top-16 upregulated genes were related to the processes highlighted by the overall enrichment analysis in this cluster, except for the *igfbp6b* whose upregulation alone points to growth suppression – a process otherwise more prominent in the downregulation cluster. Upregulation of stress protection mechanisms is reflected by increased expression of *hmox1a* that plays a role in protection against oxidative damage, which indicates that stationary-phase cells could be experiencing oxidative stress resulting from buildup of metabolic waste (36, 63). Increased expression of *rcn3* involved in maintenance of endoplasmic reticulum function and protein folding could confer protection against endoplasmic reticulum stress (64), while upregulation of *sacs2*, which is a chaperone involved in mitochondrial function and cytoskeletal organization, suggests the need to confer protection to mitochondria in order to maintain a basal level of energy production (65). Upregulation of *dap1b* known as negative regulatory of autophagy, may seem counterintuitive (66, 67). However, since the activity of this protein is chiefly regulated through phosphorylation, changes in its abundance may have a less significant effect or even indicate a compensatory response (68, 69). Similarly, upregulation of *sypl1* known to be involved in vesicle trafficking could be indicative of increased protein secretion and changes in secretory phenotype in growth-arrested cells, but could also be a compensatory response, since this gene is also known to promote cell survival in tumors by inhibiting apoptosis (70). Lastly, upregulation of numerous collagens (*col2a1b* and *col6a2* among the top-16 and several others present among the upregulated DEPs) along with lumican (*lum*), known to be involved in organization of collagen fibers, and *sparc* (osteonectin), known to modulate cell-matrix interactions, point to structural reinforcement and massive remodeling of extracellular matrix (71–74). Most of the cellular machinery processes found to change between the exponential and stationary growth phases in PAC2 were similar to those reported by other studies looking at similar growth transitions in cultured cells (52, 53).

### Tissue-specific genes significantly changing upon transition from exponential to stationary phase

Apart from the more general cellular machinery responses described in the previous section, PAC2 cells in the stationary phase also showed several responses which could be viewed as more tissue-specific. This included enrichment of exocrine system development terms in the downregulation cluster and of terms related to neuronal processes and immune-system related responses in the upregulation cluster.

Specifically, upregulated DEPs showed enrichment for biological processes “axon guidance” and “regulation of neuron projection development”. These processes were not significantly enriched in upregulated DEGs but appeared again in the multiomics list of shared upregulated DEPs and DEGs (Fig. 5). Notably, both these processes were also enriched in the full proteome, while another related term, “regulation of axonogenesis”, was highlighted in the full transcriptome annotation (Fig. 10). These observations raised the question of whether this cell line starts exhibiting neuronal characteristics. Such specialization is in principle possible, since PAC2 are derived from 24h-old zebrafish embryos and thus could retain some specific differentiation capacity (34, 75). However, many fish cell lines are in general known to have stem cell potential, so this observation may not be a specific feature of embryonic cells only. For example, strong expression of a stem cell marker *lgr5* has been observed in permanent cell lines derived from rainbow trout gut (76). Nonetheless, although our present evidence does not allow for concluding that PAC2 cells could transform into neuronal cells over time, future research could investigate their differentiation potential in this regard.

Enrichment of the terms “negative regulation of immune system process” and “immune system process” was observed in the upregulated DEGs and in the shared upregulated DEPs/DEGs list, respectively. While no significant enrichment was found among upregulated DEPs, a number of immune system-related genes were found among the top-16 genes upregulated on both the transcript and protein levels, as well as in the DEG list overall. For example, *dhx58* is a pattern recognition receptor involved in antiviral innate immunity (77), while *stat2* is a central activator of antiviral-like gene expression cascades, but is also known to play a role in regulating inflammation and promoting cell survival (78–81). *Ifi44g, ifit16 and isg15* represent downstream targets of interferon signaling, with the two former ones being interferon-induced proteins which have been linked to viral defense and inflammatory pathways and the latter one (*isg15)* being ubiquitin-like modifier which conjugates to proteins in response to interferon signaling (82). The expression of various immune system components in fish cell lines and their ability to mount active responses to immune challenges has been demonstrated previously, for example, in rainbow trout gut cells (RTgut-GC) treated with lipopolysaccharide (13) or in zebrafish embryonic line Z428 challenged by Singapore Grouper Iridovirus (42). The upregulation of these genes in the stationary phase of unchallenged PAC2 cells could point to the induction of interferon-like signaling as part of an antiviral-like response pathway, which could represent a generalized defense mechanism that can be triggered by low-level stress even in sterile cell culture condition.

### Functional characteristics of PAC2 cells revealed by multiomics profiling

To evaluate the functional capacities of PAC2 cells based on the expressed gene repertoire, we next explored the obtained proteomics and transcriptomics profiles in their entirety. Our PAC2 datasets contain almost twice as many transcripts measured as protein groups (Fig. 7). This was expected since RNAseq technology is known to provide deeper coverage than mass spectrometry-based global proteomics. However, as our analyses revealed, the level of proteomics coverage attained in the present study may already be sufficient for many research applications. Indeed, as shown in Fig. 10 and Table 3, the major functional categories represented by identified genes were shared between both datasets, and the number of associated terms was also comparable, except in the categories “Regulatory processes” and “Response to external stimuli”, where transcriptomics data achieved a two-times higher term representation (Table 3). However, higher representation of transcripts of genes with regulatory function is in fact to be expected, because many regulatory components are typically expressed at low levels and often transiently, which makes their detection on protein level even less likely. Thus, both the global proteomics and transcriptomics data obtained in our study were able to reveal highly similar biological processes despite the difference in total number of genes detected by each omics technology.

Interestingly, proteomics analysis also identified ca. 800 proteins which did not have a corresponding transcript detected in the matched samples, which showed enrichment of terms related, e.g., to extracellular space and activity of membrane receptors and ligand-gated channels. Many of these proteins appear to be membrane proteins, detected thanks to the SDS-based cell lysis procedure employed in our sample preparation protocol, which is known to efficiently solubilize many membrane proteins (49). The absence of corresponding transcripts’ detection could be due to the existing of alternative splicing isoforms and difficulties with detecting polyadenylated regions of some mRNAs (83) or to generally low transcript expression in conjunction with higher protein stability, as for example some membrane proteins with slower turnover rates could be retained even when corresponding mRNA expression has already ceased (84).

Another interesting observation was that a large group of terms highlighted in both transcriptomics and proteomics datasets was related to developmental processes and growth, such as “embryo development”, “chordate embryonic development” as well as development of “circulatory system”, “hemopoiesis”, “myeloid cell differentiation”, “connective tissue” and “cranial skeletal system”. In addition, “locomotory behavior” and “digestive tract development” were highlighted in the proteomics dataset only. This evidence reveals that some of the molecular characteristics of the PAC2 cell line could be retained from the early zebrafish embryonic tissue it was originally derived from (10, 33).

### Expression of cell morphology-related molecular markers in PAC2 cells

Molecular profiling can also help shed more light on the question of morphological classification of the PAC2 cell line. Originally, it was classified as “fibroblast-like” (33). However, the visual appearance of the PAC2 cells currently cultured in our laboratory may question this classification, as the cells exhibit rather an epithelial-like morphology (Figure 1) and express a number of proteins representative of both morphologies, as we highlighted in a short conference report published last year based on preliminary analysis of PD1 (85). The analysis of PD1/PD2 proteomics along with the transcriptomics data obtained in the current study has now confirmed the expression of both types of markers in PAC2 on both the protein and transcript level (Table 4). Namely, fibroblast-like markers expressed in PAC2 cells include fibronectin, N-cadherin and collagen type I, while expressed epithelial-like markers include E-cadherin, occludin, zona occludens 2, mucin-5AC, and syndecan (Fig. S7). Importantly, vimentin, a well-known marker of fibroblast morphology, was not detected on the protein level, while on the transcript level it was detected only in the unfiltered counts dataset and hence its expression is negligible. Comprehensive multiomics evidence obtained in our study thus reinforces the possible need to reconsider the morphological classification of the PAC2 cell line that we voiced previously (85).

### Expression of toxicologically relevant gene groups in PAC2 cells

As shown in Table 4, zebrafish PAC2 cells express a number of toxicologically relevant gene families and pathways, underscoring the relevance of this cell model for use in toxicological studies. The biotransformation capacity of cell models is frequently a matter of concern, as the extent of *in vitro* cultured cells’ ability to biotransform xenobiotics is often unclear. In PAC2 cells, for example, expression of *cyp1a* was neither detected in our study nor could be detected or induced (as checked on mRNA level) by others, in line with their observation that PAC2 cells are not able to biotransform polycyclic aromatic hydrocarbons such as BaP (86). However, we found that PAC2 cells do express aryl hydrocarbon receptor 2 (*ahr2*) and aryl hydrocarbon nuclear translocator 2 (*arnt2*), detected on both the mRNA and protein level, along with a low mRNA level of PXR, as well as *cyp1b1,* a number of *cyp2* isoforms and several other *cyp* genes (Table 4), also reported by (86). This suggests that the ability to biotransform other hydrocarbons or compounds such as estradiol could still be retained in the PAC2 cells. However, this would need to be confirmed experimentally in future studies. *Cyp2* family genes expressed in PAC2 cells (*cyp2aa1*, *cyp2aa11*, *cyp2aa6*, *cyp2ae1* and *cyp2v2*) could exhibit analogous activity to mammalian CYP2 enzymes, which are known to process diverse xenobiotics including drugs, pesticides and industrial chemicals. Further, *cyp3a65* is a xenobiotic-inducible zebrafish homolog of CYP3A (87, 88). In mammals, CYP3A is a major drug-processing enzyme known to biotransform pharmaceuticals and dietary xenobiotics (37, 89). In addition, expression of *cyp4t8* could confer the ability to process lipophilic xenobiotics, as in mammals some CYP4 isoforms, especially CYP41B, are known to biotransform some industrial chemicals, in addition to their primary internal function in fatty acid oxidation (90, 91). With regard to phase II biotransformation, PAC2 cells were already demonstrated to express a wide repertoire of GST enzymes (21), many of which are relevant for handling xenobiotics (37, 92). Multiomics evidence obtained in the current study also demonstrates the expression of UGTs, SULTs, and NATs (Table 4). Members of the *ugt1* and *ugt5* families are broadly active toward phenolic and steroid substrates and in zebrafish are known to glucuronidate bisphenol A, alkylphenol, diclofenac, and steroids (93). For members of *ugt2* family, substrate specificity is less defined, and they could be involved in internal metabolic functions instead. *Sult1* and *sult3* family members can catalyze sulfation of phenols and amine (94). Lastly, N-acetyltransferase (NAT) enzymes can be involved in detoxification of hydrazines and aromatic amines, such as beta-naphthylamine (95). PAC2 cells were also found to express a wide variety of ABC transporters, of which many support cellular defense against chemical exposure (e.g., *abcc1*, *abcc3*, *abcc4*, *abcc5, abcb6*, *abcb7*, *abcg2c*) (96–100). Further, PAC2 cells express an enormous variety of SLCs, some of which are known to respond to changes in cellular homeostasis, such as *slc30a5* (101), and *slc36a4* (37), as well as genes involved in xenobiotic transport, including family members of *slc16*, *slc22*, *slc47* (102–105).

Next to biotransformation capacity, the ability of cultured cells to mount specific molecular signaling pathways in response to chemical exposure is also an important determinant of its suitability for certain toxicological investigations. In this regard, it is interesting to note that PAC2 cells might likely detect and respond to estrogenic compounds through estrogen receptors *esr2a* and *esr2b*. These cells also express both the glucocorticoid receptor (*nr3c1*, detected on both the mRNA and protein level) and mineralocorticoid receptor (*nr3c2*, detected on mRNA level only), which could make them a useful model for studies on effects of synthetic corticosteroids. Further, PAC2 cells might also have capacity to respond to lipid-regulating and peroxisome proliferator compounds, such as fibrates and industrial chemicals like phthalates, due to them expressing several peroxisome-proliferating receptors (*pparab*, *pparda*, *ppargc1a*, *ppargc1b*), as well as liver X receptor alpha (LXRalpha, *nr1h3*) and farnesoid X receptor (FXR, *nr1h4*), known to be involved in cholesterol transport, lipid and glucose metabolism, and inflammation. The presence of PPARs, LXRalpha and FXR suggests that PAC2 cells could offer a useful model for studying metabolic disruption, especially chemical effects on lipid metabolism (106–110). We also observed, albeit at low levels, the mRNA expression of several genes from the Rev-Erb family (*nr1d1*, *nr1d2a*, *nr1d4b*), which function as circadian modulators of metabolism (111).

Additionally, we detected the expression of several clock-controlled genes, e.g., from *smad* and *tgf* families (Table 4), thus corroborating an earlier study in zebrafish larvae (112). Hence, it might be interesting to continue exploring whether PAC2 cells exhibit differential sensitivity to chemicals dependent on the daytime, as this could establish them as a model for studying interaction of metabolic rhythms with responses to xenobiotics. Lastly, PAC2 were observed to express a number of other nuclear receptors, some of which are known to act as modulators of retinoic acid signaling (e.g., nr2 family genes (113), or co-repressors of other NRs and could thus exert indirect influence on biotransformation of or response to xenobiotics in PAC2 cells.

Overall, our analysis shows that PAC2 cells express a broad molecular repertoire that includes major components of xenobiotic biotransformation across all biotransformation phases, along with a versatile suite of toxicologically relevant pathways and numerous transcriptional regulators capable of sensing a wide range of chemical stimuli and mounting transcriptional and functional responses to them. PAC2 cells could thus offer a versatile and biologically informative *in vitro* system for chemical screening and studying molecular mechanisms of toxicity. However, it should be emphasized that for a significant proportion of the genes discussed, expression was detected only on the mRNA level (Table 4). Hence, further confirmation of the protein-level expression, along with functional validation, would be needed to confirm the presence and activity of the corresponding enzymes, receptors, and response pathways, which should be addressed in the future studies.

## Conclusions

This study presents a comprehensive molecular profiling of the zebrafish PAC2 cell line, combining an extensive global proteomics analysis across cell population growth phases complemented with a matched transcriptomics dataset at two timepoints corresponding to exponential and stationary phases. Proteomics analysis of two independently generated sample series delivered comparable results despite significant differences in sample preparation protocols, which confirmed the high robustness of the employed DIA-based global proteomics workflow. Furthermore, despite the differences in the total number of gene identifications, both omics approaches highlighted similar biological categories and processes, indicating that the proteome coverage obtained was sufficiently deep to reveal meaningful functional insights. This study thus paves the way for a broader application of state-of-the-art global proteomics techniques for molecular characterization of fish cell models. Multiomics analysis revealed that PAC2 cells express a broad array of toxicologically relevant genes and pathways, including, e.g., xenobiotic biotransformation enzymes, stress response pathways, and a suite of chemical-responsive nuclear receptors. These findings reinforce the suitability of PAC2 cells as a versatile and well-characterized *in vitro* model for in-depth studies of fish cell biology, which can utilize different gene networks expressed at proliferative and stationary cell population growth phases, as well as omics-enhanced exploration of chemical toxicity mechanisms, further leveraging the availability of well-developed molecular annotation for this species. Beyond toxicology, molecular characterization data generated in this study can be further explored to evaluate PAC2 model usefulness for other disciplines, including fish virology, immunology, and aquaculture (e.g., for studying disease mechanisms and host responses to pathogens). Moreover, the analysis pipelines and approaches developed in this work will provide a blueprint for molecular baseline characterization of other fish cell lines, particularly for a suite of cell lines derived from different organs of rainbow trout, foreseen to constitute a vital component of the fish invitrome for animal-free toxicity prediction. This work thus strengthens the mechanistic knowledge base supporting the use of fish cell lines in aquatic toxicity testing and risk assessment.

## Materials and methods

### PAC2 cell culture and growth curve characterization

The permanent zebrafish embryonic cell line PAC2 (Cellosaurus accession number: CVCL_5853) was routinely cultured at 26°C ± 1°C in T-75 flasks (Thermo Fisher Scientific) with 10 mL of Lebovitz’s cell culture medium (L-15; PAN-Biotech), supplemented with 5% fetal calf serum (FCS; Eurobio Scientific). Sub-culturing was performed after reaching full confluence and involved washing with Versene (Gibco™), detachment with trypsin (0.25% in Dulbecco’s phosphate buffered saline; PAN-Biotech), cell counting by Casy cell counter (OLS OMNI Life Science, Bremen, Germany) and seeding at one to two million into a new T-75 flask in fresh culture medium. Cells were routinely observed using a microscope (Nikon Eclipse TS2).

### Growth curves in 96-well plates - experimental design overview

To characterize cell population growth phases, PAC2 cells were seeded in the 96-well plates (Greiner Bio-One CELLSTAR 96 Well, transparent, F-bottom, with lid) at a density of 3500 cells in 200 µL/well of the L-15 medium supplemented with 5% FBS; empty wells were loaded with 200 µL culture media. The plates were sealed with transparent sterile adhesive sheets (Thermo Scientific), the lid was sealed with Parafilm® and the cells were cultured at 26°C ± 1°C. Seven days after seeding, the old culture medium was removed by inverting the plate and gently tapping it onto a clean tissue paper. Fresh medium was then added at 200 µL/well. Cell viability and cell counting data were acquired at multiple timepoints (days) post-seeding from different wells of the same 96-well plate. In most experiments, 12 wells (rows B to G in columns 8 and 10) were seeded with cells: the top six were assigned for cell viability and the bottom six for cell counting. For the cell viability assays, six adjacent wells without cells served as blanks. The cell viability assays were performed using the fluorescent dyes alamarBlue™ (Invitrogen, indicator of metabolic activity) and 5-carboxyfluorescein diacetate, acetoxymethyl ester (CFDA-AM, Invitrogen, indicator of membrane integrity), according to previously established protocols (18, 35, 114). After dye addition and 30 min incubation in the dark at 26°C ± 1°C, fluorescence intensity was measured on a Tecan Infinite M200 plate reader with the following parameters: bottom well reading; 530nm excitation and 595nm emission for alamarBlue™; 493nm excitation and 541nm emission for CFDA-AM. Cell count data was acquired on the Biotek Cytation 5 (Agilent) using the nuclear stain Hoechst 33342 (Invitrogen), according to the previously described procedures (115), using standardized settings for the DAPI channel (excitation 377 nm, emission 477 nm) and Gen5 software (version 3.10.06). All measurement data were exported to Excel for further analysis. Fluorescent intensity readings from cell-containing wells were corrected by subtracting the average fluorescent intensity readings of the cell-free wells (blanks). Growth curves were plotted in RStudio (version 4.4.2) with ggplot2’s geom_smooth (span = 1), using the LOESS (locally estimated scatterplot smoothing) function.

### Sample collection for omics analyses

Based on the growth curve information, collection timepoints were set at days 3, 8, and 14 post-seeding, corresponding to the early exponential, mid-exponential, and stationary cell culture growth phases. To ensure sufficient amounts of protein and RNA material, cell culturing for omics analyses was performed in 6-well plates seeded at a density of 90000 cells in 4mL/well culture medium. Relative to the well surface area, this cell density is slightly lower than the one used for cell seeding in the growth curve experiments in the 96-well plates described above; it was selected based on several scaling experiments of different cell seeding densities between the two plate formats (data not shown). Plates were sealed and incubated in the dark at 26°C ± 1°C. Culture medium was exchanged after 7 days post-seeding. On days 8 and 14, corresponding to exponential and stationary growth phases, two sets of samples, one for global proteomics and one for transcriptomics, were collected from the same plate (top row was used to collect cells for the transcriptomics samples and bottom row for the proteomics samples), thus ensuring direct pairing of multiomics samples. In addition, day 3 samples were collected for proteomics only. These experiments were performed in four biological replicates with different cell passage numbers (P85-88), and experiments started on different days. Thus-collected samples underwent proteomics and transcriptomics analyses as described below, generating proteomics dataset 1 (PD1) and transcriptomics dataset (TD). In addition, a second set of samples collected at days 4, 7, 11, 18 and 28 post-seeding was included for global proteomics analysis in our study, generating proteomics dataset 2 (PD2). These samples, representing days 4, 7, 11, 18, and 28 post-seeding, originated from another PAC2 cell culture growth experiment reported in (35), performed in three biological replicates. In contrast to PD1 samples which were trypsin-digested and desalted (see below) prior to mass spectrometry analysis, PD2 samples comprised a non-bound fraction collected during Fe-NTA phosphopeptide enrichment procedure, described in (35). Further differences between the two proteomics sample sets are described in Table 1 above.

### Global proteomics analysis: sample collection and preparation for MS analysis

For proteomics sample collection, cells were washed three times with ice cold 1x PBS (without Ca^2+^ and Mg^2+^) before adding lysis buffer containing 5% sodium dodecyl sulfate (SDS) and 5% triethylammonium bicarbonate buffer (TEAB) in nanopure water. Cellular lysis was facilitated using scrapers with swiveling 13 mm blades. Lysed cells were pooled from 3-6 wells (depending on the time point / amount of biomass per well) in a 1.5 mL Protein LoBind® Eppendorf tube, which was then heated for 10 min at 90°C with a ThermoMixer (Eppendorf), cooled to room temperature and stored at -70°C until protein extraction. Both PD1 and PD2 samples underwent tryptic digestion following the same procedures based on S-trap™ (Protifi) Micro high recovery protocol (HaileMariam et al., 2018). Briefly, frozen samples were brought to room temperature and sonicated for 5 min in a bath sonicator to facilitate recovery of wall-adsorbed proteins. After this, the lysate was passed 6-10 times through a syringe needle (Sterican, 21G 0.80 x 40mm) to shear the DNA. Protein concentration in the lysates was determined using the Pierce™ BCA Protein Assay Kits (Thermo Scientific™) with absorbance measured on Tecan (Infinite M200) plate reader at 562 nm wavelength. Tryptic digest for PD1 samples started with 50 µg (day 3) or 100 μg protein (day 8 and 14) and for PD2 with 200 µg for all samples. Disulphide bond reduction and alkylation were successively performed for 30 min in the dark each, using, respectively, 5 mM tris(2-carboxyethyl)phosphine–hydrochloride (TCEP) added at volume ratio 1:50 and 25 mM of the alkylator 2-iodoacetamide (IAA) at volume ratio 1:20. Next, the lysate mixture was acidified with formic acid (FA; 100%, final pH <1) and thoroughly mixed by repeated pipetting. Binding/wash buffer (10% TEAB of pH adjusted to pH 7.55 with FA and 90% methanol) was then added to each sample at volume ratio 6:1 and mixed by pipetting before transferring the sample mixture to the S-Trap micro column placed in a 2 mL receiver tube. The S-Traps were centrifuged at 4000 g for 30 s to trap proteins, the flow-through was reloaded onto the trap, and the procedure was repeated twice. After this, 200 µl of binding/wash buffer was added, centrifuged at 4,000 g for 30 s, and the flow-through was discarded. This procedure was repeated two more times. Finally, the S-Trap column was centrifuged (1 min, 4000g) to fully remove the binding/wash buffer and then transferred to a new 1.5 mL Eppendorf tube. To each S-Trap, 120 µl of the digestion buffer containing elution buffer 1 (50mM TEAB in nanopure water), trypsin (1:100 *w/w*; Roche), and 0.5% 0.5M CaCl2, was added to each S-Trap, spun through, re-loaded and incubated for 12-14 h at 37°C. The next day, elution buffer 1 was added and centrifuged for 1 min at 4000 g. This step was repeated with elution buffer 2 (0.2% FA in nanopure water) and elution buffer 3 (50% acetonitrile in nanopure water). All eluates of the same sample were collected in one tube, centrifuged, and dried in the vacuum concentrator (Eppendorf). After this step, the handling of PD1 and PD2 samples started to differ. PD1 samples were resuspended in buffer A (0.1% FA in nanopure water), desalted on the tC18 cartridges (Sep-Pak, Vac 1cc (50mg), #WAT054960), eluted with buffer B (0.1% FA and 80% acetonitrile in nanopure water), centrifuged, dried, resuspended in buffer A, filtered with 0.45 µm centrifugal filter (Amicon Ultrafree MC), transferred into glass AS-vials with a 0.1 ml insert and stored at 4°C until MS measurement. PD2 samples underwent the Fe^3+^-NTA phosphopeptide enrichment procedure described in (35) and comprise the non-bound fractions containing non-phosphorylated peptides which were collected during the washing steps in this procedure, pooled, dried and then treated similarly to PD1 samples after final drying step.

### Global proteomics analysis: data acquisition and processing

Peptides (1-4 µg, see Table 1) were separated on a nanoLC system (Dionex Ultimate 300 UHPLC, Thermo Scientific) using an in-house made C18 reverse-phase analytical column (25 cm x 75 µm ID) with self-pulled tip, packed with ReproSil-Pur, 120 Å pore size, C18-AQ, 3 µm bead size (Dr. Maisch GmbH, Germany), and coupled with a Nanospray Flex Series ion source NG^TM^ to the Orbitrap Fusion^TM^ Lumos^TM^ Tribrid^TM^ (Thermo Scientific) mass spectrometer operated in data-independent acquisition mode (DIA) over 138 min gradient at a flow rate of 300 nL/min, including 92 min linear gradient from 2% to 35% eluent B (0.2% FA, 2% nanopure water, 97.8% acetonitrile). Source parameters were 2400 V for positive ion spray and 275°C ion transfer tube temperature. MS was set for expected LC peak width of 60 s with a default charge state of 2. Full scan set at 120000 resolution, scan range of 350-1150 m/z, normalized AGC target 200% and maximum injection time 100 ms; DIA was performed using 41 variable-width isolation windows. MS^n^ experiment was set at HCD collision energy of 28%, 30000 resolution, scan range of 200 – 1800 m/z, AGC target 400%, maximum injection time 54 ms. Most samples were run with two technical replicates, except PD1-day 3 samples (three technical replicates) and the third biological replicate of PD2 (one technical replicate). Raw LC-MS/MS data were converted with HTRMS converter (version 18.2) and processed in Spectronaut® 18.0 Pulsar (version 18.2.230802.50606, Biognosys AG, Schlieren, Switzerland) (40). Proteins were inferred based on peptide fragment ions identified using directDIA, a method embedded in Spectronaut that uses a FASTA sequence database instead of a spectral library. Search FASTAs included reference proteome of zebrafish (Danio rerio, UP000000437) downloaded from UniProt on 19.9.2023 (46695 entries) and NIHMS1873978_supplement_contaminant (381 entries) (116). Trypsin was selected as a digestion enzyme with one miscleavage allowed, carbamidomethylation was set as fixed modification, and methionine oxidation and N-terminal acetylation as variable modifications. False discovery rate (FDR) was set to 0.01 and quantification was performed on MS2 level. Identified proteins were combined into protein groups where peptide information did not allow their unambiguous distinction from each other, and final quantification was performed on the level of protein group using only protein group-specific peptides. Normalization was performed locally using sparse profiles and imputation was disabled. Following the initial processing in Spectronaut, data was exported and further analyzed using Excel, RStudio and Perseus (version 2.0.11.0) (117). The mass spectrometry proteomics data has been deposited to the ProteomeXchange Consortium via the PRIDE (118) partner repository with the dataset identifier PXD066565.

### Global proteomics: downstream data analysis

For each biological replicate, technical replicates (if any) were averaged and results (i.e., one value per protein group, per biological replicate) used for protein identification counts, where each protein required at least one measured value in one of the biological replicates for the respective condition to be counted as having identified that protein. For downstream quantitative analysis, proteins which were not present in at least 70% of the replicates of the entire dataset were removed. Values were then log2 transformed and missing values were imputed based on the Perseus default settings, where missing values are randomly assigned a value artificially generated from a set of low values mimicking the parametric distribution of the real values per sample (see Fig. S2 for the distribution of real and imputed data in every biological replicate). Data centering was then performed by subtracting the median of the sample from every value within the corresponding sample. Next, a PCA was performed using the *prcomp* function, with the first two components visualized using *ggplot2*. Samples were grouped by day, with convex hulls (*chull*) to illustrate group boundaries and biological replicates labeled using *ggrepel*. Variance explained by each component was indicated on the axes. Differential expression analysis was conducted in Perseus separately for each dataset. PD1 and PD2 were analyzed by a one-way analysis of variance (ANOVA), while an unpaired Student’s t-test was used on a subset of PD1 (days 8 and 14) matched with TD. In both tests, significance was assessed by permutation-based false discovery rate (FDR) approach, with a threshold of FDR<0.05 and 250 randomizations. No grouping was preserved during permutation. Following statistical testing, differentially expressed protein groups (DEPs) were z-score normalized by subtracting the mean and dividing by the standard deviation of protein group intensities across conditions. Hierarchical clustering was then carried out using Euclidean distance and complete linkage. Heatmaps of the clustered DEPs were generated in RStudio using the *pheatmap* package for visualization. To identify the top regulated DEPs, a volcano plot was generated using visualization tools from Perseus.

### Transcriptomics analysis: sample collection and library preparation

To collect the samples for transcriptomics analysis, cells were washed with ice cold 1x PBS before adding RNeasy Plus lysis buffer (QIAGEN, Ref. 1053393). Cellular lysis was facilitated using scrapers with swiveling 13mm blades. Lysed cells were harvested from multiple wells in 1.5 mL DNase/RNase-free Eppendorf Collection Tubes (QIAGEN) placed on ice, vortexed at maximal speed for 1 min and immediately stored at –70°C until further processing. RNA extraction was performed according to manufacturer’s instructions using the RNeasy Plus Mini Kit (QIAGEN, Ref. 172034266), including a gDNA removal step. RNA quality was analyzed with the RNA ScreenTape Assay for TapeStation 4200 System (Agilent), following the assay instructions and using the electronic ladder. For all samples, the measured RNA Integrity Number (RIN) was equal to 10. After this quality control, the samples were transported in dry ice to the Genomic Diversity Centre (GDC), ETH Zürich, which provides access to managed instruments and expertise in genetic techniques and genomic data. Here, samples were treated with DNase using the DNase I Amplification Grade Kit (Invitrogen, Ref. 18068015) to eliminate any residual gDNA, using 1 µg of RNA for each sample and following the kit instructions. After DNAse treatment, three random samples were tested again with the TapeStation to ensure the quality of RNA was not changed. All tested samples showed RINs equaling 10.

To prepare the mRNA library, DNAse-treated RNA samples were first quantified using the Qubit RNA Broad Range (BR) Assay Kit (Invitrogen, Ref. Q10211) following kit instruction. Measurements were performed in a 384-well black plate (Faust, Ref. 4000246) using a plate reader Spark M10 (Tecan). mRNA library was prepared with starting amount of 0.5 µg for each sample, using the TruSeq Stranded mRNA Library Preparation Kit from Illumina (96 Samples, Ref. 2002055), which includes poly-A mRNA enrichment through oligo(dT) magnetic beads. For increased demultiplexing accuracy and reduced sample misassignment during sequencing, IDT for Illumina – TruSeq RNA Unique Dual (UD) Indexes v2 (96 Indexes, 96 Samples, Ref. 20040871) were used. The whole procedure was performed following manufacturer’s instructions but using half-volumes of all reagents. This modification of the protocol was previously tested and validated (internal trial; data not shown). Following the preparation, the concentration of each mRNA library was measured using a Qubit dsDNA BR Assay Kit (Invitrogen, Ref. Q32853) following the kit instructions. One sample (day 14, biological replicate 3) appeared to be lost during library preparation and was therefore not included in downstream analyses. The remaining mRNA libraries were normalized and pooled following the specifications required by the sequencing platform.

After pooling, residual adapter dimers were removed by an additional mRNA library purification step using AMPure XP beads (self-prepared by the GDC) using a bead-to-sample volume ratio of 0.95:1 and following the mRNA library purification steps described in the TruSeq Stranded mRNA Library Preparation protocol. The samples were then air-dried for 15 min, removed from the magnetic stand, and gently resuspended in 70 µL of nuclease-free ultrapure water. After 2 min incubation, samples were centrifuged at 280 rcf for 1 min and placed back into the magnetic rack. Once the liquid became clear, the supernatant containing the cleaned mRNA library was collected into a new tube. The final quality check of the pooled, cleaned library was performed using the D1000 ScreenTape Assay for TapeStation Systems (Agilent).

### Transcriptomics analysis: sequencing and data processing

The pooled mRNA library was sequenced by the Functional Genomics Center of University of Zurich and ETH Zurich (FGCZ). Before the main sequencing, mRNA quality check was performed with iSeq (Illumina). This additional step of short, low-output sequencing provided information about the optimal loading concentration, library quality, and index balance. The index balance for under-represented samples detected by the iSeq analysis was adjusted by respective spiking into the final mRNA library, followed by evaluation of sample concentration and quality of sample using the ScreenTape assay. The final sequencing was performed on an Illumina NovaSeq X system using a 300-cycle 10B flow cell and yielded an average of approximately 20 million paired-end reads (2 × 150 bp) per sample. Raw data (sample reads provided by the sequencing facility (FGCZ)) were pre-processed by the GDC, including the pseudo-alignment which was performed using Kallisto (version v0.50.1) (119) and the reference transcriptome from Ensembl (*Danio rerio*, GRCz11/GCA_000002035.4). Transcripts remapping was on average above 83%, with most samples yielding between 12-33 million aligned reads. The fragment length ranged between 168-209 bp. The RNAseq transcriptomics data has been deposited to the Gene Expression Omnibus (GEO) (120) with the dataset identifier GSE303229.

### Transcriptomics analysis: downstream data analysis

Transcript-level abundance estimates from Kallisto (in HDF5 format) were imported into RStudio (version 4.4.2) and summarized to the gene level counts based on the reference transcriptome. Raw counts per million (CPM) values were extracted and normalized using transcript length and effective library size, thus accounting for the sequencing depth and gene length bias. The normalized dataset was log-transformed. Differentially expressed genes (DEGs) analysis was performed using the *edgeR* package (version 4.6.2) (121). A design matrix accounting for the two experimental conditions of interest (exponential vs. stationary growth phases) was generated. Genes with low expression were filtered using the *filterByExpr* function. The filtering step removed approximately 13000 genes, going from 30628 to 17570 genes. The *glmQLFit* function, which applies a negative binomial model and robust estimation to reduce the impact of outliers, was used to model the variation in gene expression across samples. Differential expressions were tested applying a quasi-likelihood F-Test (*glmQLFTest* function) and corrected by FDR<0.05 obtained using the Benjamini-Hochberg (BH) adjustment. Following statistical testing, DEGs were z-score normalized and hierarchically clustered using Euclidean distance and complete linkage. Heatmaps of clustered DEGs were generated in RStudio using the *pheatmap* package for visualization. To visualize the data, transcripts were filtered according to the transcript per million (TPM) value, where each transcript’s geometric mean was calculated across samples and within each cell population growth phase. Normalized data with TPM>1 was selected for PCA, performed using the *prcomp* function with default parameters. The first two components were visualized using *ggplot2* (v3.5.2). Samples were grouped by day, with convex hulls (*chull*) to illustrate group boundaries. Variance explained by each component was indicated on the axes. To identify the top regulated DEGs, a volcano plot was generated using visualization tools from Perseus.

### Multiomics data integration and functional annotation analysis

Identifiers (UniProt accessions for PDs and Ensembl IDs for TD) were converted to official gene symbols using DAVID Bioinformatics (122, 123), which were then used for overlap analysis (Venn diagrams) and functional annotation analysis. Transcripts used in analysis were filtered according to the TPM value, where each transcript’s geometric mean was calculated across samples and within growth phase, and transcripts with TPM<1 were excluded from the overlap analysis. Venn diagrams were analyzed using the InteractiVenn webpage (124) and re-drawn using BioRender.com. Functional annotation and enrichment analysis for Gene Ontology (GO) biological process (BP; GOTERM_BP_FAT), molecular function (MF; GOTERM_BP_FAT), cellular component (CC; GOTERM_MF_FAT) and KEGG pathway (KEGG) terms were performed using DAVID Bioinformatics and respective gene symbol lists. When analyzing the complete list of identified protein groups and transcripts, no background was applied. When analyzing DEPs and DEGs, the full PAC2 proteome and transcriptome, respectively, were used as background references. Statistically significant GO BP terms (FDR<0.05) identified in the previous steps were clustered based on semantic similarity (default settings, specifically medium/0.7 resulting list and SimRel) using the *Danio rerio* species background in Revigo (version 1.8.1) webpage (125). The process included removal of obsolete terms. To represent the functions of the entire proteome and transcriptome, the Interactive Graph results from Revigo were exported in an XGMML file format and transferred to Cytoscape version 3.10.3 (126). In Cytoscape, terms (represented by circles) were color-coded according to pre-established cellular functional categories based on the proximity and nomenclature of the circles (i.e., common words used throughout circles found in close distance from each other). To represent the functions of up- and down-regulated DEPs and DEGs, resulting GO terms and KEGG pathways were grouped according to gene similarity using DAVID Bioinformatics in-build clustering algorithm. Terms and pathways of interest were selected and represented using horizontal lollipop charts inspired from the design of the visualization tools used in interactive web-based platform, ShinyGO 0.81 (127).

### Generation of multiomics data model

A multiomic model was built with RStudio (v4.5.1) for the matched PD1 and TD samples (day 8 and day 14) using normalized proteomics and transcriptomics data, created as described in the corresponding sections above. A horizontal (shared sample model) multiblock data integration partial least-square (PLS) model was implemented using the *block.pls* function from the mixomics R library (v.3.21) (128). PLS maximizes the covariance between datasets (blocks) and generates latent variables that capture their shared variations. Canonical method and near-zero variance filtering were used as arguments. PLS plot capturing variance per biological replicate and per dataset was generated using the *plotArrow* function.

## AI statement

Several RStudio scripts used for figure preparation were refined with the assistance of ChatGPT (OpenAI).

## Author contributions

M.-O.D.: conceptualization, formal analysis, investigation, validation, visualization, writing – original draft preparation, writing – review & editing; J.B.: formal analysis, investigation, validation, writing – review & editing; N.W.: investigation, resources, writing – review & editing; D.L.R.: formal analysis, visualization, writing – review & editing; M.R. – writing – review & editing; R.S. – investigation, methodology, writing – review & editing; C.v.B.: conceptualization, funding acquisition, writing – review & editing; K.S. – conceptualization, funding acquisition, project administration, resources, writing – review & editing; K.G. – conceptualization, formal analysis, funding acquisition, investigation, methodology, project administration, resources, supervision, validation, visualization, writing – original draft preparation, writing – review & editing.

## Acknowledgements

This research was funded by the Swiss National Science Foundation through National Research Programme 79 “Advancing 3R – Animals, research and society”, project number 407940_206439, “Expanding the fish invitrome towards a modular, socio-technical framework for animal-free prediction of chemical toxicity to fish”. Transcriptomics data presented in this paper were produced and analyzed in collaboration with the Genetic Diversity Centre (GDC), ETH Zurich.

## Notes

### Competing Interest Statement

The authors have declared no competing interest.

