## supplementary figures and files for "Multiomics profiling of zebrafish embryonic cell line PAC2 across growth phases to assess its relevance for toxicological studies": PAC2_supplementary_figures_subm250726.pdf


By

Mihai-Ovidiu Degeratu[1,2], Jessica Bertoli[1,2], Nikolai Huwa[1], David Lopez Rodriguez[1,3,4], Marion Revel[1], René Schönenberger[1], Colette vom Berg[1], Kristin Schirmer[1,2], Ksenia Groh[1*]


**List of Figures**


**Figure S1.** Growth curve of the PAC2 cell line based on membrane integrity measurements.

**Figure S2.** Histograms of the biological replicates from proteomics datasets.

**Figure S3.** Absolute abundance differences of protein groups shared between proteomics datasets 1 and 2 (PD1 and PD2).

**Figure S4.** Impact of peptide loading concentration in LC-MS/MS runs on correlation between average protein group abundancies in proteomics datasets 1 and 2 (PD1 and PD2).

**Figure S5.** Principal component analysis (PCA) of individual proteomics datasets.

**Figure S6.** Co-expression plots of the z-score normalized differentially expressed protein groups (DEPs) identified as significantly down- or upregulated in proteomics datasets 1 (PD1) and 2 (PD2).

**Figure S7.** Expression levels of selected markers of fibroblastic and epithelial cell types.

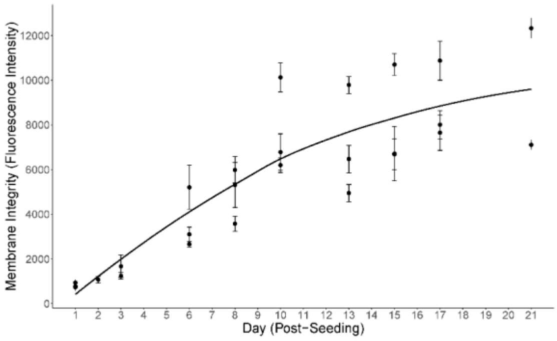


**Figure S1. Growth curve of the PAC2 cell line based on membrane integrity measurements**. PAC2 cells were seeded in 96-well plates and cultured for three weeks. Cell culture medium was exchanged once on day 7 post-seeding. Membrane integrity based on 5-caroxyfluorescein diacetate, acetoxymethyl ester (CFDA-AM) fluorescence was recorded in three biological replicates at multiple timepoints (days) throughout the culturing duration. Data are shown as mean ± SD of technical replicates (n=3-12) for each biological replicate. Trend line was added using LOESS smoothing (geom_smooth, span = 1) to visualize overall changes over time; no further statistical modeling was applied.

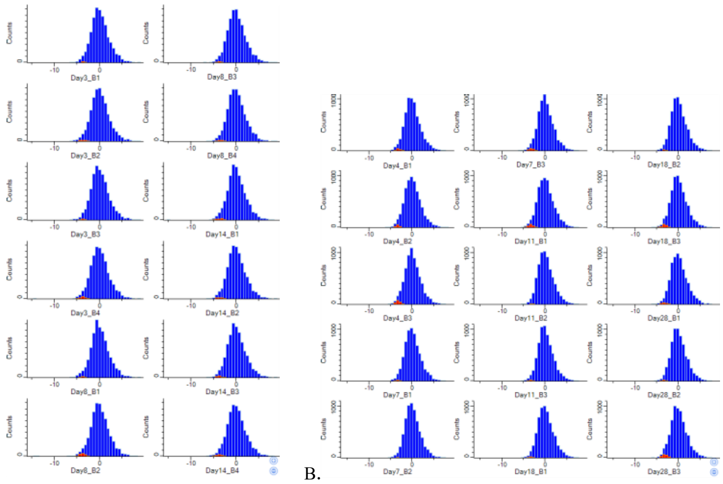


**Figure S2. Histograms of the biological replicates from proteomics datasets.** Histograms from proteomics dataset 1 (PD1) and 2 (PD2) are shown in panels A and B, respectively. Protein group abundance values were log2-transformed and filtered to retain only those with values in at least 70% of biological replicates. Remaining missing values were imputed (shown in the red bins, located left of 0 on x-axis) and centred to 0 (by subtracting the median per sample). Biological replicates are labelled as B1 to B4 for PD1 and B1 to B3 for PD2.

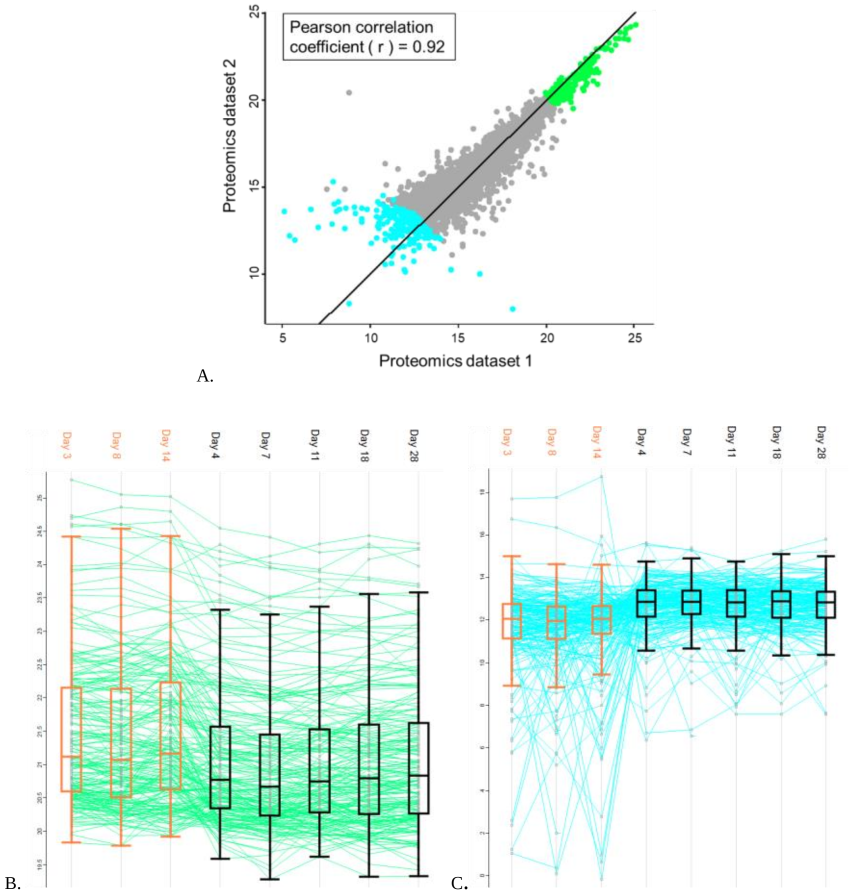


**Figure S3. Absolute abundance differences of protein groups shared between proteomics datasets 1 and 2 (PD1 and PD2).** (A) Correlation and density plot of 6803 protein groups (subset with >70% detection) shared between PD1 and PD2. For each dataset, abundance values of each protein group were averaged across biological replicates from all conditions (Days). The diagonal line indicates equality (x = y); points above the line have higher values in PD2, and those below have higher values in PD1. Protein groups are color-coded according to abundance: green represents the 200 most abundant and blue the 200 least abundant, based on the average values across all biological replicates and conditions from both datasets. (B–C) Profile plots of the top 200 most abundant (B) and least abundant (C) protein groups shown in (A), with box plots illustrating the distribution of abundance values across conditions (Days) for each dataset (orange for PD1; black for PD2).

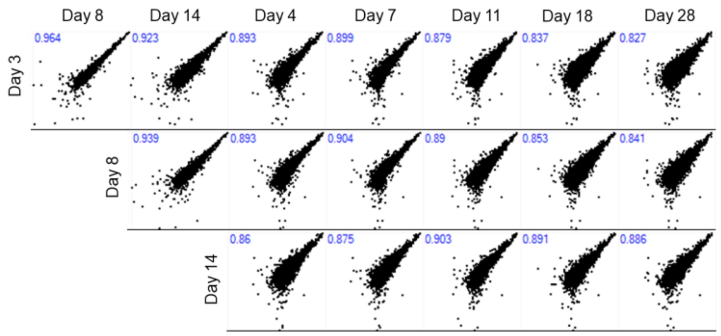


**Figure S4. Impact of peptide loading concentration in LC-MS/MS runs on correlation between average protein group abundances across different samples in proteomics datasets 1 and 2 (PD1 and PD2)**. Correlation plots show protein group abundances (averaged across biological replicates) between conditions: day 3 (1 µg on-column loading per sample), day 8 and day 14 (2 µg on-column loading per sample) from PD1; and days 4, 7, 11, 18, and 28 (4 µg on-column loading per sample) from PD2. Pearson correlation coefficients are shown in each plot.

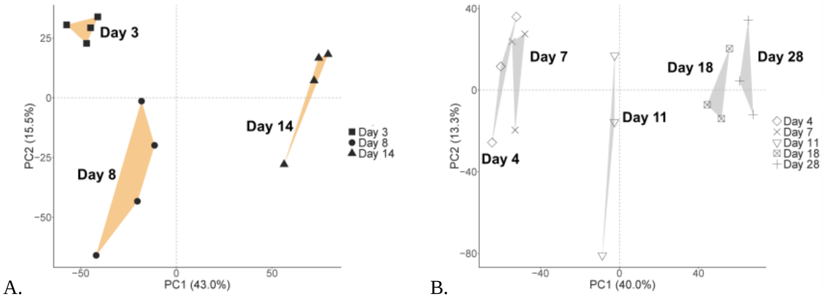


A. B.

**Figure S5. Principal component analysis (PCA) of individual proteomics datasets**. PCA plots are shown for proteomics dataset 1 (PD1; orange; panel A) and proteomics dataset 2 (PD2; grey; panel B). PCA was performed on filtered, $log_2$-transformed, and zero-centred data, which comprised 6751 protein groups for PD1 and 6927 protein groups for PD2. Convex hulls (orange and grey areas for PD1 and PD2, respectively) show the outer boundaries formed by replicates within each group.

**Commented [GK1]:** @Mihai: can you please also do the same plots for Day 8 and Day 14? It's actually interesting to see if 2 ug compared to 4 ug would show a better correlation than 1 ug. And, after thinking about this more, I think we do have SOME effect of peptide loading amounts. Else how would you explain that it did get lower (<0.9) when comparing with 4 ug? What would also be interesting to see is, how will the PD2 samples compare to each other (will the r be higher?) Actually, we should be expecting SOME deviation from line of unity, because samples also differ between each other (different growth phases) and so we expect them to have differential protein expression. Question is, how much. Can you do a figure that shows correlation for Day 3 on one line, followed by Day 8 on the second line (starting under Day 14), then for Day 14 (starting under day 4) and so on? And day8-day28 should be indicated on top, while comparators (day 3 -day 18) on the left side.

**Commented [DM2R1]:** Done

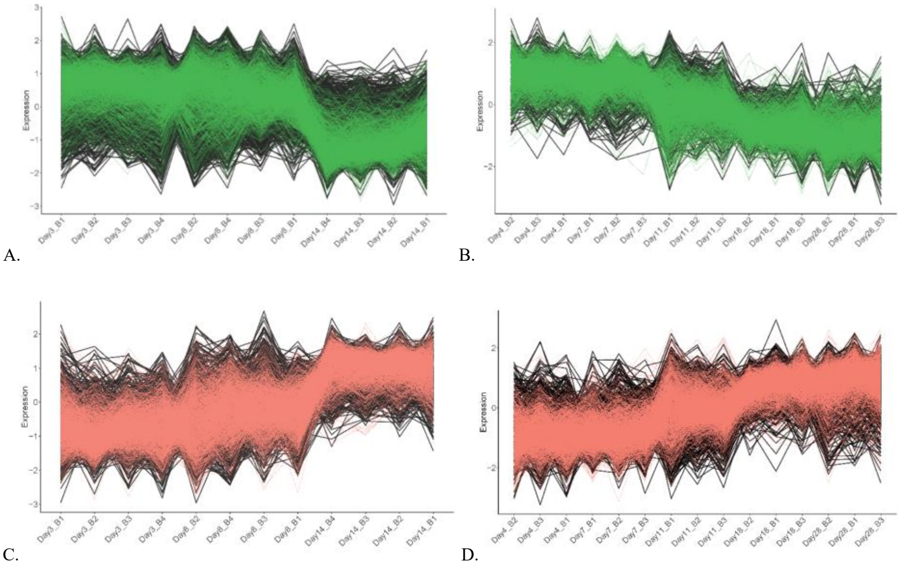


**Figure S6. Co-expression plots of the z-score normalized differentially expressed protein groups (DEPs) identified as down- or upregulated in proteomics datasets 1 and 2 (PD1 and PD2)**. In each plot, all lines represent the total number of differentially expressed proteins (DEPs) in the corresponding case, while coloured lines represent the DEPs shared between PD1 and PD2. Panels (A) and (B) show downregulation clusters in PD1 (2317 DEPs) and PD2 (1622 DEPs), respectively, with 1247 shared DEPs (green lines). Panels (C) and (D) show upregulation clusters in PD1 (2133 DEPs) and PD2 (1909 DEPs), respectively, with 1273 shared DEPs (pink lines). Abbreviations: B, biological replicate.

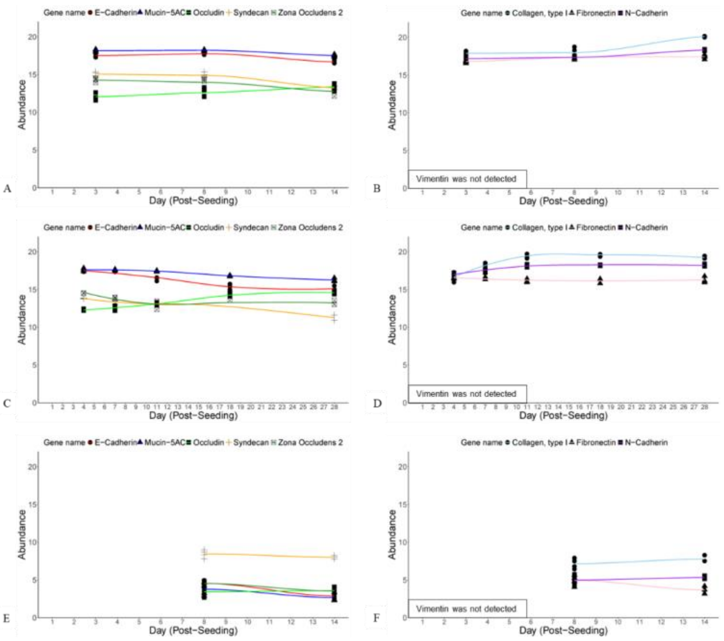


**Figure S7. Expression levels of selected markers of fibroblastic and epithelial cell types**. Markers were selected to represent epithelial-like morphologies (panels A, C, and E) and fibroblast-like morphologies (panels B, D, and F). Expression data are shown for selected markers using proteomics dataset 1 (PD1; panels A and B), proteomics dataset 2 (PD2; panels C and D), and the transcriptomics dataset (TD; panels E and F). For PD1 and PD2, protein group abundance values were used; for TD, transcript levels were represented as TPM values. All data were $\log_2$-transformed and visualized in RStudio. Epithelial markers full name, including gene symbol and UniProt accession: cadherin 1, type 1, E-cadherin (epithelial) (*cdh1*, Q90Z37); mucin-5AC (*prr36b,* A0A8N7UTE4); occludin (si:ch73-61d6.3, A0A8M1PUX9); syndecan (*sdc2*, Q66I66); zona occludens 2 (*tjp2b*, Q6DHS7). Fibroblastic markers full name, including gene symbol and UniProt accession: vimentin (*vim*, A0A8M2BL67); collagen, type I (*col1a1b,* A0A8M2B227); fibronectin (*fn1a*, B0S602); cadherin 2, type 1, N-cadherin (neuronal) (*cdh2*, Q90275).
